# Transcription Factor Localization Dynamics and DNA Binding Drive Distinct Promoter Interpretations

**DOI:** 10.1101/2022.08.30.505887

**Authors:** Kieran Sweeney, Megan N. McClean

## Abstract

Environmental information may be encoded in the temporal dynamics of transcription factor (TF) activation and subsequently decoded by gene promoters to enact stimulus-specific gene expression programs. Previous studies of this behavior focused on the encoding and decoding of information in TF nuclear localization dynamics, yet cells control the activity of TFs in myriad ways, including by regulating their ability to bind DNA. Here, we use light-controlled mutants of the yeast TF Msn2 as a model system to investigate how promoter decoding of TF localization dynamics is affected by changes in the ability of the TF to bind DNA. We find that yeast promoters directly decode the light-controlled localization dynamics of Msn2 and that the effects of changing Msn2 affinity on that decoding behavior are highly promoter dependent, illustrating how cells could potentially regulate TF localization dynamics and DNA binding in concert for improved control of gene expression.

## INTRODUCTION

To survive changes in their environment, cells transmit environmental information through signaling pathways to transcription factors (TFs), which bind DNA and regulate the gene expression response. Signaling pathways often exhibit a bowtie topology where multiple environmental signals converge on a single TF (Csete and Doyle, 2004). In these cases, how does the TF activate the appropriate set of genes needed for each environmental signal? One way cells may overcome this challenge is by encoding additional environmental information in the temporal dynamics of TF activation. For example, extracellular calcium causes the *Saccharomyces cerevisiae* TF Crz1 to translocate to the nucleus in one of two modes, continuous or pulsatile, and recent work has shown that Crz1 target genes decode its localization dynamics by preferentially activating in response to one mode over the other (Cai et al., 2008; Chen et al., 2020). At least 10 yeast TFs and a variety of mammalian TFs exhibit similar stimulus-specific dynamics (Batchelor et al., 2011; Dalal et al., 2014; Tay et al., 2010; Yissachar et al., 2013). In addition to regulating TF localization dynamics, cells possess other mechanisms for modulating TF activity, each offering the opportunity to encode information. The mammalian inflammatory response TF NF-κB exhibits stimulus specific dynamics—it pulses in and out of the nucleus in response to tumor necrosis factor-α (TNFα), but undergoes sustained nuclear localization in response to bacterial lipopolysaccharides (LPS)—and is subject to post-translational modifications (PTMs) that regulate its ability to bind DNA (Chen et al., 2002; Crawley et al., 2013; Ea and Baltimore, 2009; Kiernan et al., 2003; Werner et al., 2005). Similarly, gamma irradiation causes short bursts of the tumor suppressor p53 that are sufficient to activate DNA repair genes, while activation of apoptotic genes may both involve sustained p53 activity and PTMs that improve its ability to bind DNA targets (Batchelor et al., 2011; Gu and Roeder, 1997; He et al., 2019; Purvis et al., 2012; Sykes et al., 2006; Vonderach et al., 2019).

A prime example of bowtie topology is the yeast TF Msn2, a C2H2 zinc finger protein (the largest structural class of TF in eukaryotes) that regulates over 200 stress defense genes. Multiple signaling pathways (PKA, TOR, SNF) converge on Msn2, which in turn plays a key role in regulating the cellular response to a variety of environmental stresses (Gasch et al., 2000; Petrenko et al., 2013). Under normal growth conditions, Msn2 is phosphorylated by protein kinase A (PKA) and resides primarily in the cytoplasm, but following environmental stress, Msn2 is dephosphorylated and translocates to the nucleus where it regulates its target genes by binding stress response elements (STREs) in their promoters (Görner et al., 1998). The identity and magnitude of the environmental stresses is encoded at least in part by the nuclear localization dynamics of Msn2: hyperosmotic shock causes an early, continuous pulse of Msn2 nuclear localization whose duration is dose-dependent, while glucose starvation causes a similar early pulse that’s followed by short, sporadic bursts of nuclear localization whose frequency is dose-dependent (Hao and O’Shea, 2012). These stresses elicit distinct transcriptional responses and previous work using chemical inhibition of PKA to control Msn2 localization showed that target genes decode Msn2 dynamics by exhibiting differential responses to the amplitude, duration, and frequency of Msn2 nuclear localization (Hansen and O’Shea, 2013, 2015a; Petrenko et al., 2013).

We therefore use Msn2 as a model system to investigate the interplay of TF localization dynamics and binding affinity in gene induction. We partially disconnect Msn2 from upstream regulation of its localization, which we instead control with light. By combining optogenetic control of Msn2 and high-throughput microscopy, we probe the relationship between Msn2 nuclear localization dynamics and the expression of target genes. We quantify the signal decoding behavior of these genes using a computational model, which suggests that changing Msn2 affinity would have highly promoter dependent effects on decoding behavior. We test this prediction by exploiting known mutations to the Msn2 DNA binding domain (DBD) to create high and low affinity light-controlled Msn2 mutants and perform additional optogenetic experiments to quantify how such changes affect promoter decoding of Msn2 localization dynamics. By combining experiments with computational models and sensitivity analysis, we identify properties of a promoter that allow it to exhibit differential responses to high and low affinity Msn2 mutants. We also measure the effect of tuning TF affinity on gene expression responses to natural stimuli. Lastly, by controlling Msn2 independently of upstream regulation, we find evidence that nuclear translocation triggers the degradation of Msn2, which has a substantial effect on the expression of certain promoters. This study contributes a fundamental understanding of how TF affinity and localization dynamics operate together to control gene expression.

## RESULTS

### Construction and optimization of a light-controlled Msn2

To control Msn2 without perturbing its upstream regulators, we used CLASP (Chen et al., 2020) to directly control the nuclear translocation of Msn2 with light (**Figure 1A**). In this optogenetic system, Msn2 is fused at the N-terminus to Zdk1, a peptide that preferentially binds a plasma membrane anchor in the dark, and at the C-terminus to mScarlet and yeLANS, which features a *Avena sativa* LOV2 domain with a Jα tail bearing a light activated nuclear localization signal (NLS) that is largely inaccessible to the nuclear import machinery in the dark. When excited by blue light, Zdk1 undocks from its plasma membrane anchor and yeLANS undergoes a conformational change that exposes its NLS to the nuclear import machinery, causing Msn2-CLASP to be imported into the nucleus where it can activate target genes (**Figure 1B**). When the blue light is turned off, Msn2-CLASP is rapidly exported from the nucleus due a constitutive nuclear export signal (NES) within yeLANS. Since strong blue light doses have been reported to induce a stress response in yeast involving Msn2 (Bodvard et al., 2017), we also created Msn2-dCLASP (deactivated CLASP) controls lacking Zdk1 and yeLANS to verify that nuclear localization events were strictly due to optogenetic control.

**Figure 1.**
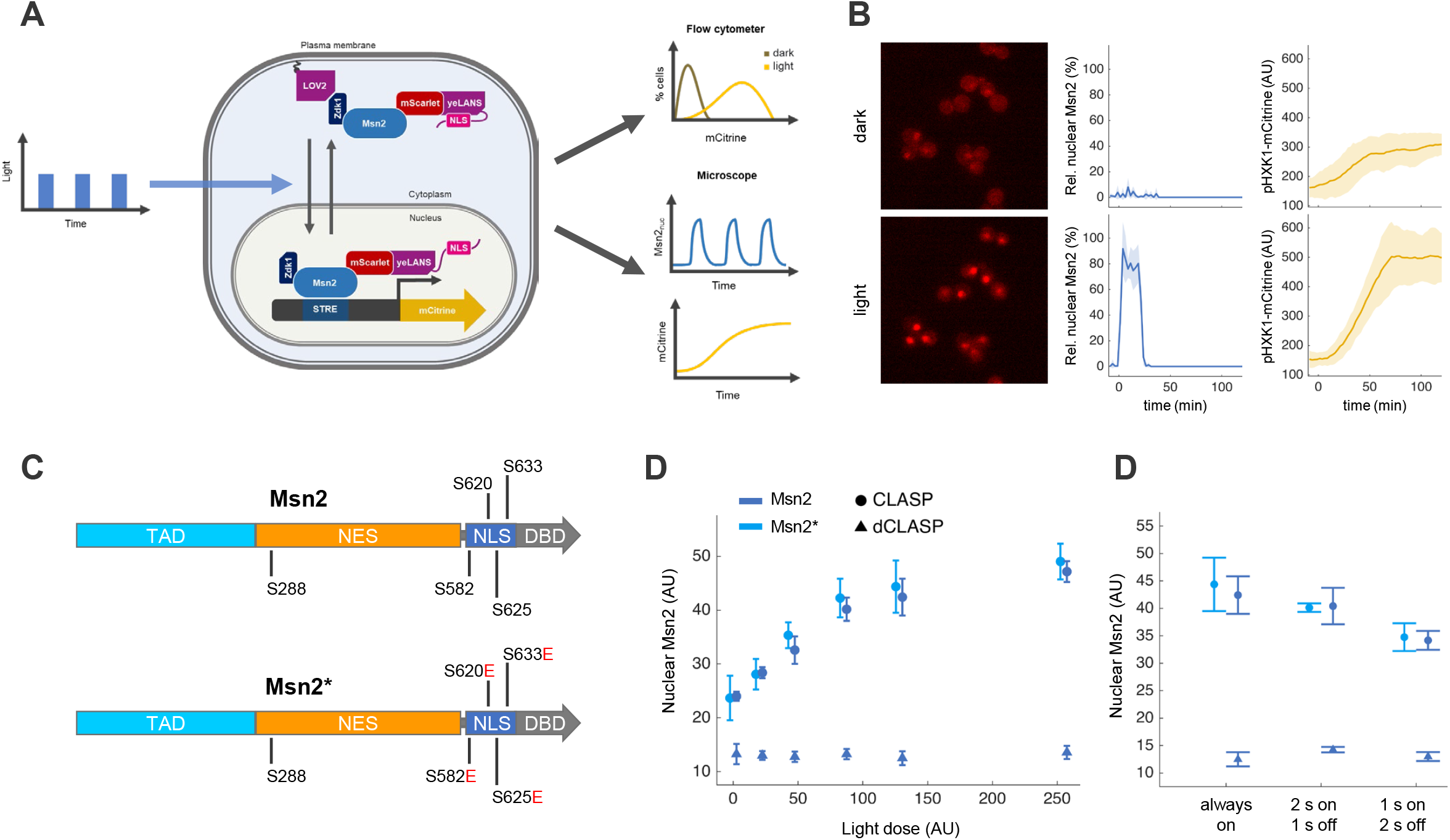
Optogenetic control of Msn2 localization using CLASP A. A schematic of the Msn2-CLASP system and experiments in which time varying light doses drove corresponding patterns of Msn2-CLASP nuclear localization and gene expression that were measured either by flow cytometry or fluorescence microscopy. B. (Left panels) Fluorescence microscopy images showing Msn2-CLASP localizing to the nucleus in response to 255 AU blue light. In the dark, Msn2-CLASP was primarily localized to the cytoplasm, but could stochastically pulse into the nucleus. Msn2-CLASP localization (middle panels) and expression of a p*HXK1*-mCitrine reporter (right panels) in the dark and in response to a 10 min pulse of 128 AU blue light (indicated by blue shading). Solid lines and shaded regions respectively represent the mean and standard deviation of Msn2-CLASP localization or reporter expression for three biological replicates, each with at least 47 cells. C.Schematic showing relevant functional domains of Msn2—the transactivation domain (TAD), nuclear export signal (NES), nuclear localization signal (NLS), and zinc finger DNA binding domain (DBD)—and residues that were mutated to identify Msn2 mutants more suitable for optogenetic control. Msn2* has a wild type (WT) TAD, NES, and DBD but features four serine to glutamic acid mutations in its NLS that inhibit stress-regulated nuclear localization. D. (Left panel) Absolute nuclear levels of Msn2-CLASP, Msn2*-CLASP, and Msn2-dCLASP were measured in response to 15 min of constant blue light with intensities ranging from 0 – 255 AU. These light doses correspond to irradiances of 0 – 405 μW/cm^2^ (**Figure S1C**) (Right panel) Nuclear Msn2-CLASP and Msn2*-CLASP in response to 128 AU blue light with varying degrees of PWM. Points and error bars represent the mean nuclear Msn2 level of three biological replicates, each with at least 43 cells. All measurements were acquired by fluorescence microscopy.

However, Msn2 stochastically pulses in and out of the nucleus in the absence of stress (Gasch et al., 2017), which caused it to weakly activate some target genes in the dark, even when tagged with CLASP (**Figure 1B**). To disconnect Msn2 from upstream regulation of its localization, we made phosphomimetic mutations to key PKA-regulated phosphosites (**Figure 1C**) in the NLS and NES of Msn2 that control the activity of these domains (Hao et al., 2013). We mixed the mutated domains in a modular fashion and tagged the resulting Msn2 mutants with CLASP. We screened the resulting Msn2-CLASP mutants for low basal nuclear localization, reduced stochastic pulsing, and the ability to localize to the nucleus and activate target genes in light (**Figures S1A – S1B**). We selected an Msn2 mutant (Msn2*) with four inhibitory serine to glutamic acid mutations in its NLS domain for further study. Msn2*-CLASP exhibited a graded response to light, which plateaued by about 128 AU, and reduced stochastic pulsing in the dark (**Figure 1D** and **Figures S1D – S1E**). Pulsing a 128 AU light dose on for 2 s and off for 1 s caused only a modest decrease (5 – 10%) in nuclear accumulation compared to continuous illumination, despite a 33% reduction in overall light dose (**Figure 1E**). Thus, we were able to identify the minimum light doses needed to achieve a given level of Msn2*-CLASP localization.

### Light sweep experiments reveal differential promoter responses to Msn2 localization dynamics

We next used optogenetic control of Msn2* to probe how promoters respond to defined patterns of Msn2* localization. Using an optoPlate (Bugaj and Lim, 2019) mounted over a 96 well plate on an inverted fluorescence microscope, we delivered time-varying light doses to a reporter strain and imaged the corresponding pulses of Msn2* nuclear localization and any subsequent fluorescent reporter induction (**Figure 1A and Figures S2 – S5**). Over these “light sweep” experiments, we exposed 12 reporter strains to a set of 14 light programs that drove Msn2* nuclear localization timecourses with a defined amplitude, duration, or pulsing behavior (**Figure 2A and Figure S5A – S5B**). Excluding a negative control, all reporter strains featured mCitrine expressed under a promoter with at least one STRE (**Figure S5C)**. No reporter was strongly activated by Msn2*-dCLASP following a 50 min 100% amplitude dose of blue light (**Figure S5D**), indicating that, without CLASP, Msn2* was not activated by the light doses used.

**Figure 2.**
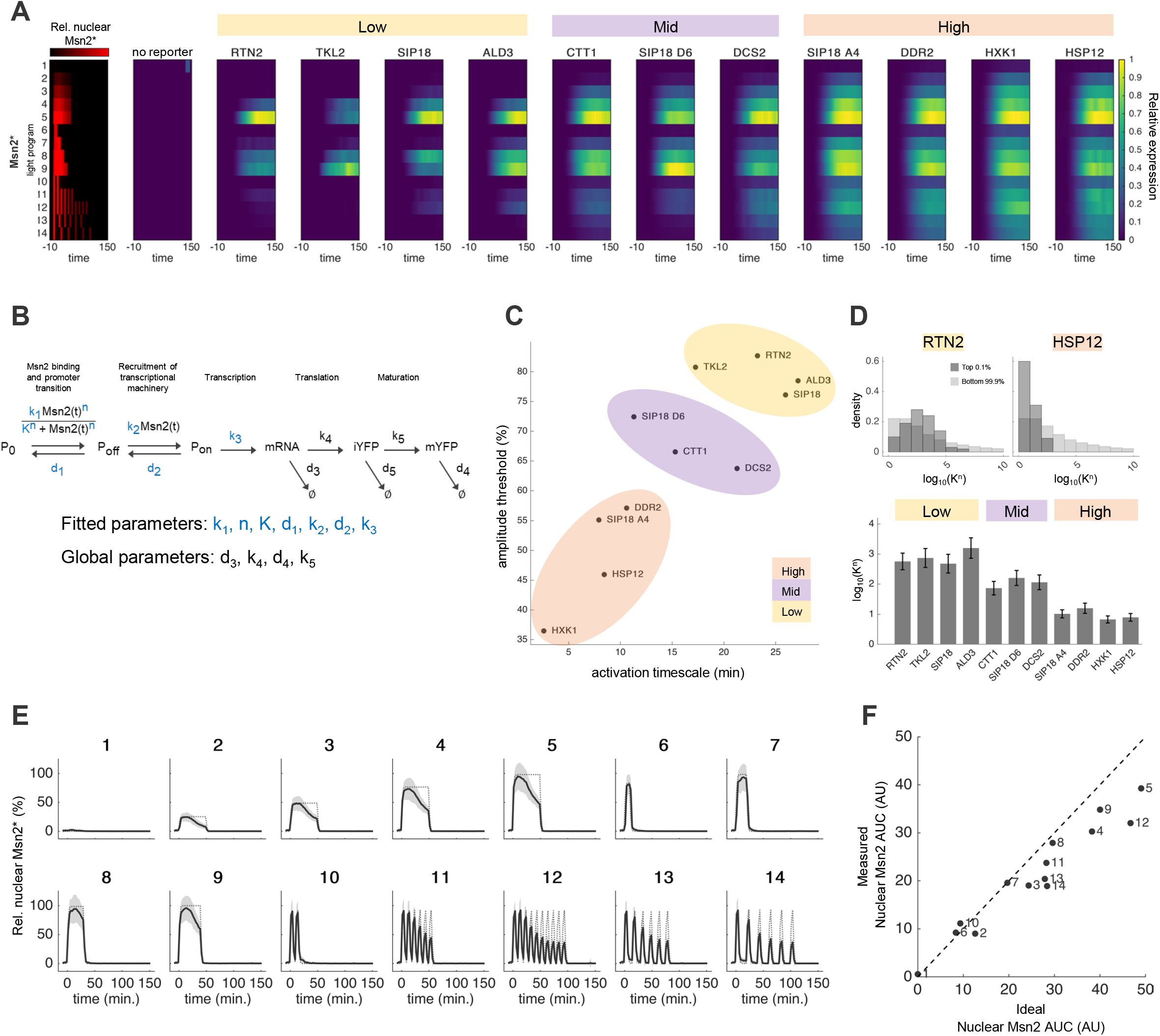
Light sweep experiments and localization triggered Msn2 degradation A.Light sweep experiments probe how promoters decode the nuclear localization dynamics of Msn2*. Each row represents a light program, which drove Msn2 localization (left) and reporter expression (right). Msn2* localization measurements were pooled over many experiments and represent thousands of cells per condition. Expression measurements were normalized to maximum expression per reporter across all conditions and represent ∼100 - 600 cells per condition from at least three biological replicates. See also **Figure S2 – S4** and **Figure S5**. B.A mathematical model of gene expression. Promoter-specific parameters (blue) were estimated by least squares fitting to the time-resolved mCitrine (YFP) measurements from all light programs. Pooled Msn2 localization measurements were used as the input Msn2(t). Global parameters (black) were taken from the literature. See also **Figures S2 – S4**. C.Categorization of promoters based on how they decoded single pulses of nuclear Msn2*. The amplitude threshold and activation timescale were calculated using the gene expression model and respectively denote the amplitude and duration of nuclear Msn2* needed to reach half the maximum promoter activity (k_3_P_on_) attained by a 50 min 100% amplitude ideal pulse of nuclear Msn2*. Promoter categories were obtained by k-means clustering of the amplitude thresholds, activation timescales, and predicted values of K^n^. D. (Top panel) Predicted values of K^n^ obtained from the gene expression model. Top 0.1% (top 100) of parameter sets for *RTN2* (dark grey) are enriched for high predicted values of K^n^ versus the bottom 99.9% of parameter sets (light grey); two-sample Kolmogorov-Smirnov (KS) tests showed that these distributions differed from each other (p = 0.0015). The top parameter sets for *HSP12* are enriched for low predicted values of K^n^ versus remaining parameter sets; two-sample KS tests showed that these distributions differed from each other (p<10^−24^). (Bottom panel) Predicted affinity of each promoter for Msn2* where bars and error bars denote the mean and 95% confidence interval of log_10_(K^n^) for the top 0.1% of parameter sets for each promoter. E.Measured versus ideal Msn2 localization timecourses showing localization triggered degradation of Msn2. Solid lines and shaded grey areas represent the mean and standard deviation of Msn2* localization measurements pooled from 79 experiments. Dashed lines represent ideal Msn2 localization (absent degradation) calculated from a computational model of Msn2 localization. F.Measured nuclear Msn2 AUC versus ideal nuclear Msn2 AUC per light program. Measured Msn2 AUC deviated from ideal Msn2 AUC most strongly for long duration light programs.

Our light sweep experiments showed that the signal decoding behavior of Msn2 target genes previously reported in response to modulation of PKA activity persists when Msn2 nuclear localization is controlled directly, without pertubring upstream signaling (**Figure 2A**). For example, *HSP12* and *HXK1* were activated by every light program and exhibited a graded response to the area under curve (AUC) of nuclear Msn2*, effectively integrating the nuclear Msn2 signal. In contrast, *RTN2* and *SIP18* were switch-like and filtered out short, low amplitude, and pulsatile pulses of nuclear Msn2*. We turned to a computational model of gene expression to quantify this behavior (**Figure 2B**). We modeled expression as a time-dependent function of nuclear Msn2 in which a promoter transitions from an initial off state (P_0_) to an intermediate off state (P_off_) to an active, transcribing state (P_on_). We pooled Msn2 localization measurements across multiple experiments, used the resulting composite signal as the model input Msn2(t), and fit the model for each promoter by identifying the parameter sets that best recapitulated the measured expression time courses across all 14 light programs (**Figure S2 – S4**). Since a range of parameter sets could predict gene expression with comparable error, we ranked the parameter sets by fit and selected the best performing ones for further analysis (**Figure S6B**).

### Gene expression model reveals high, mid, and low sensitivity promoter groups

Based on previous studies (Hansen and O’Shea, 2013, 2015a), we used the model to calculate amplitude thresholds and activation timescales, which respectively describe the amplitude and duration of a square pulse of nuclear Msn2 needed to attain half-maximum promoter activation (**Figure 2C** and **Figure S6C**). The amplitude threshold and activation timescale were linearly related (R^2^ = 0.69): promoters with low amplitude thresholds (like *HXK1, HSP12, SIP18 A4*, and *DDR2*) had short activation timescales, while promoters with high amplitude thresholds (like RNT2, *TKL2, SIP18*, and *ALD3*) had long activation timescales. Both the amplitude threshold and activation timescale were inversely related to the predicted value of K^n^ for each promoter. K^n^ captures the half-maximum point and slope of the curve relating nuclear Msn2 to the rate of transition for P_0_ to P_off_ and is inversely related to the affinity between the promoter and Msn2. The top parameter sets for promoters with high amplitude thresholds and long activation timescales, like *RTN2*, were enriched for high values of K^n^, implying a low affinity for Msn2 (**Figure 2D, top panel**). In contrast, promoters with low amplitude thresholds and short activation timescales, like *HSP12*, were enriched for low values of K^n^, implying a high affinity for Msn2. We therefore quantified the affinity of each promoter for Msn2 as the mean value of log_10_(K^n^) for the top 0.1% of parameter sets (**Figure 2D, bottom panel**).

Clustering the amplitude thresholds, activation timescales, and predicted values of K^n^ revealed three groups of promoters: high, mid, and low sensitivity. The high sensitivity promoters had low predicted values of K^n^, low amplitude thresholds, and short activation timescales, reflecting an ability to respond rapidly to small amounts of nuclear Msn2. In contrast, the low sensitivity promoters had high predicted values of K^n^, high amplitude thresholds, and long activation timescales, reflecting their tendency to filter out short, low amplitude doses of nuclear Msn2. The mid sensitivity promoters were characterized by intermediate predicted values of K^n^ and amplitude thresholds but had activation timescales that overlapped both the high and low sensitivity groups.

### Nuclear localization of Msn2 triggers a degradation process that continues outside the nucleus

Over the course of the light sweep experiments, we found decay in the Msn2 nuclear localization timecourses (**Figure 2E**) consistent with reports that nuclear accumulation and DNA binding causes Msn2 degradation (**Figure 2E and Figures S7A – S7B**) (Durchschlag et al., 2004; Lallet et al., 2004; Sadeh et al., 2011). By comparing the measured Msn2 localization time courses (**Figure 2E, solid lines**) with predicted (or “ideal”) time-courses (**Figure 2E, dotted lines**) obtained from a published model of Msn2 localization that does not incorporate degradation (Hansen and O’Shea, 2013), we found evidence that Msn2 degradation began about 14 min after Msn2 first entered the nucleus (**Figure S7C**) but continued in the cytoplasm (see **Figure S7D**).

Msn2 degradation cofounds simple comparisons between light programs, as there was a mismatch between the AUC of the measured nuclear Msn2 signal and the AUC of the ideal of nuclear Msn2 signal (**Figure 2F**). The mismatch was pronounced for long duration, pulsatile light programs: for example, following 30 min of continuous illumination (light program 8), the measured and ideal AUC were roughly equal, while six 5 min pulses of illumination separated by 15 min (light program 14) had a similar ideal AUC but a much lower measured AUC. As discussed below, such degradation may set a timer on gene expression, particularly for low sensitivity promoters.

### Changes to the DNA-binding affinity of Msn2 have the strongest effect on low sensitivity genes

TFs have a range of DNA binding affinities, but often exhibit sub-maximal affinity for their target sites (Crocker et al., 2016). Previous work showed that increasing or decreasing the DNA binding affinity of Msn2 respectively increases or decreases the expression of a target gene at steady-state (Stewart-Ornstein et al., 2013). However, our gene expression model predicted that the effects of such changes are highly promoter dependent (**Figure 3A**): they have a stronger effect on the predicted expression of low sensitivity genes like *RTN2* than on high sensitivity genes like *HSP12*. Moreover, it was unclear how changing the DNA binding affinity of a TF would affect the ability of its target promoters to decode nuclear localization dynamics.

**Figure 3.**
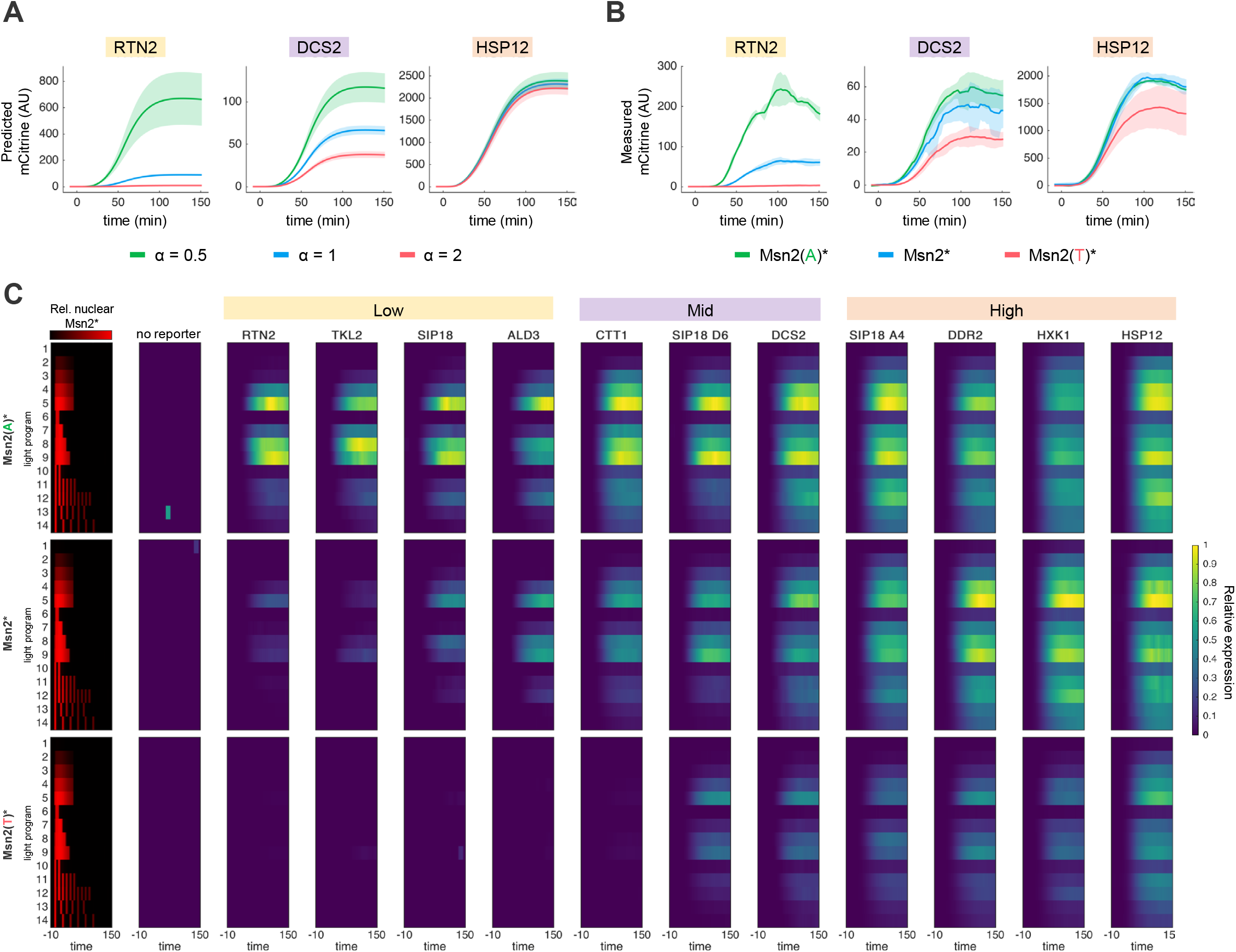
Predicted and measured expression show that changes in Msn2 affinity have a strong effect on low sensitivity promoters and a weaker effect on high sensitivity promoters A.Predicted gene expression was calculated by simulating the response of each promoter to a 50 min 100% amplitude pulse of nuclear Msn2* while scaling the K term in the gene expression model (see **Figure 2B**) by a factor α, equaling either 2, 1, or 0.5. Lines and shaded regions represent the mean and 95% confidence interval of the predicted expression for the top 10 parameter sets. B.Measured expression following a 50 min 100% amplitude pulse of Msn2(A)*, Msn2*, and Msn2(T)*. Lines and shaded regions represent the mean and standard deviation of expression for three biological replicates. C.Light sweep experiments probing how promoters decode the nuclear localization dynamics of Msn2(A)*, Msn2*, and Msn2(T)*. Each row corresponds to a light program, which drove Msn2 localization (left) and subsequent gene expression (right). Msn2 localization measurements were pooled over many experiments and represent thousands of cells per condition. Gene expression measurements were normalized to the maximum expression level per reporter across all conditions and Msn2 DBD mutants. Expression measurements for each condition represent ∼100 - 600 cells from at least three biological replicates. See also **Figure S2 – S4**.

To test the model predictions, we exploited known mutations to the DBD of Msn2 (**Figure S8A**). We created Msn2(A)*, which has a high predicted affinity to its targets and is transcriptionally hyperactive (Pfanzagl et al., 2018; Reiter et al., 2013) and Msn2(T)*, which has a low predicted affinity and reduced transcriptional activity (Stewart-Ornstein et al., 2013). With CLASP, both Msn2(A)* and Msn2(T)* localized similarly to Msn2* (**Figure S6B**). To measure how changes in Msn2 affinity affected promoter decoding of its nuclear localization dynamics we performed additional light sweep experiments with Msn2(A)* and Msn2(T)* (**Figure 3B and Figure 3C**). In general, increasing Msn2 affinity increased reporter expression and reduced the amplitude threshold and activation timescale of a promoter such that lower pulse amplitudes and durations were needed to achieve a given level of promoter activity (**Figure S8B)**. In agreement with the model predictions, these changes had a strong effect on the expression of low sensitivity promoters like *RTN2* and a comparatively weak effect on the expression of promoters with a high predicted sensitivity for Msn2 like *HSP12* (**Figure 3B**).

Overall, the high sensitivity promoters were strongly induced by all three Msn2 mutants (**Figure 3C**) and differences in their expression between mutants were typically small compared to the low sensitivity promoters (**Figure 4A**). Expression of these promoters generally increased linearly with nuclear Msn2 AUC, though in some cases approached saturation at high pulse amplitudes (**Figure 4Ai and Figure 4Aii**). There was no high sensitivity promoter where expression for Msn2(A)* was significantly higher than for Msn2*, suggesting that there was little benefit to further increasing Msn2 affinity for high sensitivity promoters. In fact, doing so may have a cost, as expression of the high sensitivity genes *HXK1* and *DDR2* was weakly, but consistently lower for Msn2(A)* versus Msn2 (**Figure S4**).

**Figure 4.**
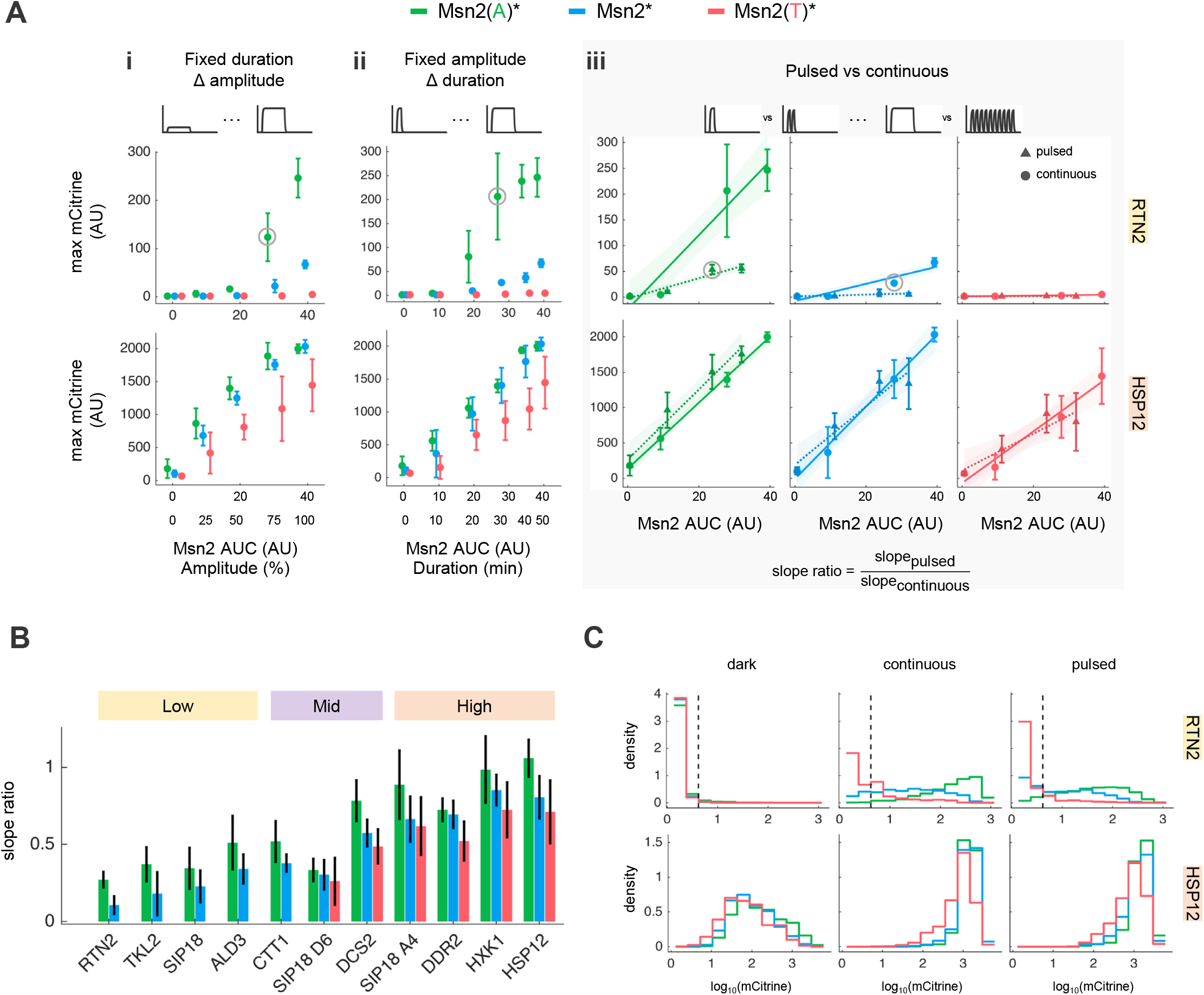
Analyzing the decoding behavior of high and low sensitivity promoters. A.Maximum reporter expression following (i) a 50 min pulse of each Msn2 DBD mutant with amplitudes ranging from 0 – 100% or (ii) a 100% amplitude pulse of nuclear localization with durations ranging from 0 – 50 min. (iii) Maximum fluorescent reporter expression in response to continuous vs pulsed doses of each Msn2 DBD mutant. Circles represent maximum expression following 0, 10, 30, or 50 min continuous pulses of 100% amplitude nuclear Msn2. Triangles represent maximum expression following 0, 2, 6, or 10 five min pulses of nuclear Msn2 with 100% amplitude and 5 min interpulse durations. In all plots, points and error bars represent mean and standard deviation of expression for three biological replicates. Solid and dashed lines show best fit lines for continuous and pulsed conditions, respectively; shaded regions show 95% confidence interval of the best fit lines. Slope ratios were calculated as the slope of the best fit line for the pulsed conditions divided by the slope of the best fit line for the continuous conditions and are shown in **Figure 4B**. Grey circles denote measurements referenced in text. B.Slope ratios were calculated as described above. Bars and error bars represent the mean and standard deviation of the slope ratio for each promoter and Msn2 DBD mutant. C.Reporter expression for cells exposed to no light, a 30 min 100% amplitude pulse of nuclear Msn2 (continuous), or six 5 min 100% amplitude pulses with 5 min interpulse durations (pulsed). Each histogram represents the distribution of single-cell fluorescence measurements of three biological replicates for a 10 min time window centered at 120 min. Dashed line represents threshold above which *RTN2* was considered active and was calculated as the 99^th^ percentile *RTN2* level for the no light condition.

In contrast, there were generally large expression differences between Msn2 mutants for the low sensitivity promoters (**Figure 3C**). Msn2(T)* failed to activate these promoters in most cells, while Msn2(A)* increased their expression versus Msn2* and effectively enabled their induction in conditions where Msn2* did not. For example, Msn2(A)* increased the maximum expression of *RTN2* by 3.7-fold (**Figure 4Ai**) and allowed its induction for pulsatile doses of nuclear localization. In fact, expression of *RTN2* was 1.9-fold higher for six 5 min pulses of Msn2(A)* than a 30 min continuous pulse of Msn2* (**circled points, Figure 4Aiii**). The relationship between expression and Msn2 AUC for low sensitivity promoters was nonlinear: they filtered out pulses of nuclear localization with low amplitudes or short durations. Activation of the low sensitivity promoters was especially dependent on the nuclear concentration of Msn2: for example, maximum *RTN2* expression (**circled points of Figure 4Ai and 4Aii**) for a 30 min 100% amplitude pulse of Msn2(A)* (AUC = 26.8) was 1.7-fold higher than for a 50 min 75% amplitude pulse (AUC = 28.2). Notably, the expression of *RTN2, SIP18*, and *TKL2* plateaued for pulses of nuclear Msn2 longer than 30 min (**Figure S8C**). This was consistent with the hypothesis that Msn2 degradation acts as a timer on the expression of low sensitivity genes as it caused nuclear Msn2 levels to drop below their high amplitude thresholds.

### Relative ability to respond to pulsed versus continuous doses of nuclear Msn2 is primarily set by the promoter

We next investigated how the response of promoters to different dynamics inputs of Msn2 nuclear localization was affected by changes in Msn2 affinity. We calculated slope ratios (Chen et al., 2020) that quantified the relative ability of each promoter to respond to pulsatile versus continuous doses of each Msn2 mutant (**Figure 4Aiii**) by fitting lines to the measurements and dividing the pulsed slopes by the continuous slopes (**Figure 4B**). A slope ratio greater than one indicates stronger promoter induction by a pulsatile dose of nuclear Msn2 than a continuous dose, while a slope ratio less than one indicates the opposite.

There were clear differences in slope ratio between promoters: the high sensitivity promoters were strongly induced by pulsatile doses of nuclear Msn2 and, accordingly, had high slope ratios—though only *HSP12* had a slope ratio above one, for Msn2(A)*—while the low sensitivity promoters were poorly activated by pulsatile doses of Msn2 and had low slope ratios (**Figure 4B**). On the other hand, differences in slope ratio between Msn2 mutants were rarely significant and likely a consequence of degradation, which disproportionally affected the pulsatile light doses (**Figure 2D**), causing nuclear Msn2 levels to drop below the elevated amplitude thresholds typically associated with a decrease in Msn2 affinity (**Figure S8B**). To compare the effects of increasing promoter affinity by adding STREs versus increasing Msn2 affinity, we analyzed the expression of *SIP18* and its mutants *SIP18 A4* and *SIP18 D6* (**Figure 4B** and **Figure S9**). *SIP18* was weakly expressed and had a low slope ratio; adding a cluster of STREs distal to its TATA box (*SIP18 D6*) moderately increased its expression but not its slope ratio, while adding a cluster of STREs proximal to its TATA box (*SIP18 A4*) increased both its expression and slope ratio. In contrast, increasing Msn2 affinity increased the expression of these promoters but not their slope ratios. Taken together, these observations indicate that the relative ability to respond to pulsed versus continuous doses of Msn2 is primarily set by the promoter rather than the TF.

### Decreasing Msn2 affinity increases gene expression noise

We also quantified differences in the cell-to-cell variability of expression, or noise, measured in the light sweep experiments (**Figure S10A**). Overall there was a negative correlation (R^2^ = 0.81) between maximum expression and noise (**Figure S10B**). Accordingly, low sensitivity promoters were much noisier than high sensitivity promoters and decreasing Msn2 affinity generally increased noise. Msn2 nuclear localization dynamics also affected expression noise, which was higher for pulsed doses of nuclear Msn2 than continuous doses. We also analyzed how Msn2 localization dynamics and affinity affected reporter expression at a single-cell level (**Figure 4C**). For example, a pulsatile dose of Msn2(T)* minimized the expression of *RTN2*, which was (weakly) expressed in just 13.8% of cells for a pulsatile dose of Msn2(T)* versus 39.0% of cells for a continuous dose of Msn2(T)*. In contrast, a pulsatile dose of Msn2(A)* moderately activated *RTN2* and a continuous dose of Msn2(A)* maximized *RTN2* expression. Meanwhile, any of these doses of Msn2(T)* or Msn2(A)* were sufficient to robustly activate *HSP12* with relatively low expression noise (**Figure S10A**). As explored further in the discussion, these measurements hint at how the concerted regulation of TF localization dynamics and affinity may facilitate improved control of expression.

### Sensitivity analysis reveals that slow activation or fast deactivation kinetics cause gene induction by high and low affinity Msn2 mutants to diverge

In previous sections of this work, we focused on how high and low sensitivity promoters responded to the nuclear localization dynamics of each Msn2 mutant because these promoters behaved consistently. In contrast, the mid sensitivity promoters exhibited a wide range of decoding behaviors for each of the Msn2 mutants, despite having similar predicted values of K^n^. This suggested that, beyond sensitivity to nuclear Msn2, other properties of a promoter may cause large differences in expression between the Msn2 affinity mutants. To identify these properties, we sought to perform a sensitivity analysis of our gene expression model. We therefore needed to incorporate affinity differences between the Msn2 mutants into the model. To capture how changes in Msn2 affinity may affect both the half-maximum point and shape of the curve relating nuclear Msn2 concentration to promoter activation, we identified factors (α and β) by which we could scale K and n to represent binding differences between the Msn2 mutants (**Figure 5A**). Since Msn2(A)* activated the most reporters in the most conditions, we did this by first parameterizing the model with the measurements for Msn2(A)* and then fitting α and β to recapitulate the expression measured for Msn2* or Msn2(T)*. We found that scaling K and n by α = 1.75 and β = 1, respectively, best approximated the expression measured for Msn2* (**Figure 5A and Figure S11**). Similarly, scaling factors of α = 1.75 and β = 1.75 approximated the expression of Msn2(T)*, though other pairs of scaling factors performed comparably. This approach assumes α = 1 and β = 1 for Msn2(A)*.

**Figure 5.**
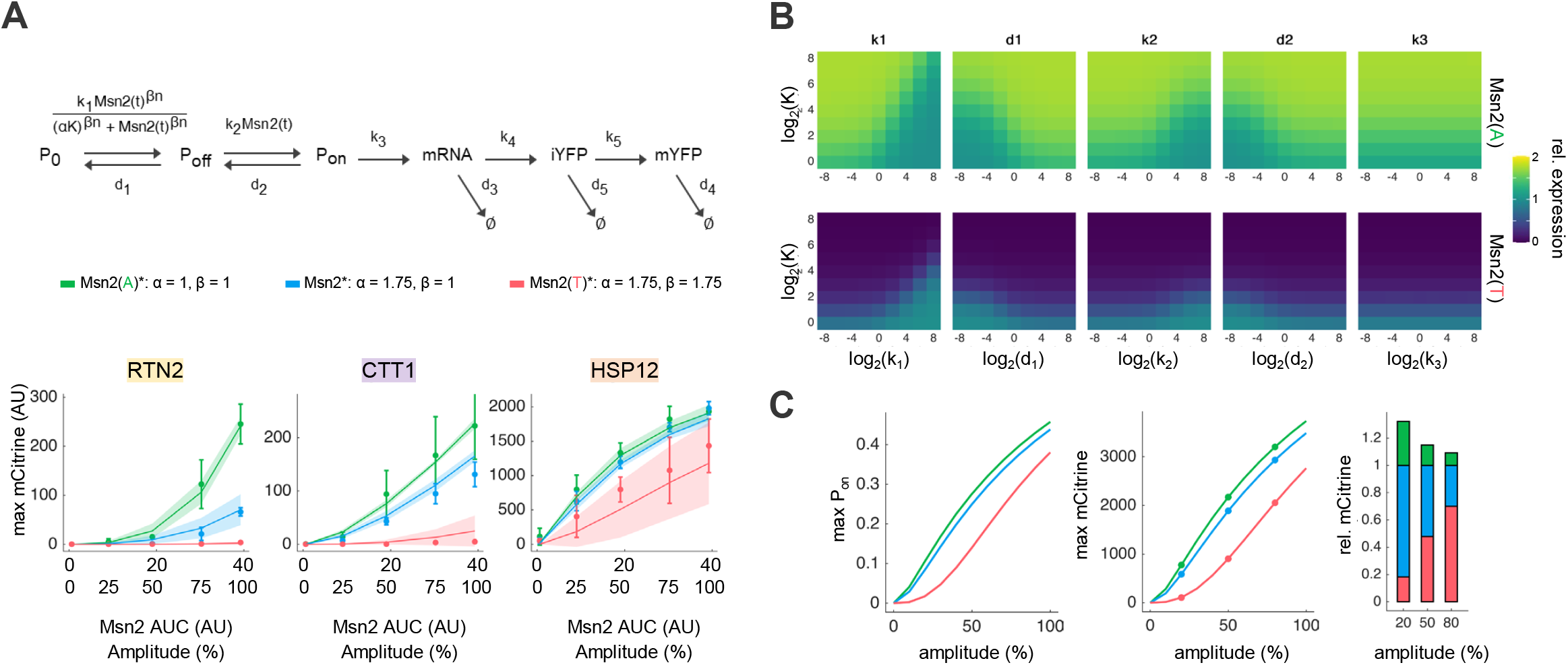
Exploring the relationship between Msn2 affinity and promoter properties using the gene expression model. A.The gene expression model was used to identify scaling factors (α and β) that represent affinity differences between Msn2 DBD mutants. Scaling factors were calculated by repeatedly simulating the expression of each reporter using the top 10 parameter sets for Msn2(A)* while scaling α and β to minimize the difference between the predicted expression and the measured expression for Msn2* or Msn2(T)* across all light programs and reporters (see also **Figure S11**). Simulated versus measured expression for the chosen scaling factors. Points and error bars show mean and standard deviation of maximum expression measured for three biological replicates of each Msn2 DBD mutant following a 50 min pulse of continuous nuclear localization with amplitudes varying from 0 – 100%. Lines and shaded regions represent mean and 95% confidence interval of expression predicted for each Msn2 DBD mutant over the same set of conditions using the top 10 parameter sets for Msn2(A)* and the indicated values of α and β. B.Simulations showing how promoter parameters affect expression for Msn2(A)* and Msn2(T)* relative to Msn2*. Expression of hypothetical promoters was simulated in response to a 50 min 100% amplitude pulse of each Msn2 DBD mutant using the values of α and β identified in **Figure 5A**. Over the simulations, each kinetic parameter (k_1_, d_1_, k_2_, d_2_, k_3_) was allowed to vary along with K (n was fixed to one). For each hypothetical promoter, relative expression was calculated by normalizing the maximum expression in response Msn2(A)* or Msn2(T)* to the maximum expression in response to Msn2*. C.Maximum P_on_ (left panel), maximum expression (middle panel), and relative expression were calculated for a hypothetical promoter (k_1_ = 1, d_1_ = 1/16, k_2_ = 1, K = 2, n = 1, d_2_ = 1, k_3_ = 1) following 50 min ideal pulses of each Msn2 DBD mutant with amplitudes ranging from 0 – 100%.

We used the best fit values of α and β to simulate the expression of hypothetical promoters for a 50 min 100% amplitude pulse of each Msn2 mutant. We individually scaled the kinetic parameters k_1_, d_1_, k_2_, d_2_, and k_3_ over a range of values of K and n and for each hypothetical promoter calculated the maximum expression for each Msn2 mutant (**Figure S12**), as well as the relative expression versus Msn2*, which we calculated by normalizing the maximum expression for each Msn2 mutant to the maximum expression for Msn2* (**Figure 5B**). As expected, decreasing promoter sensitivity (by increasing K or n) caused absolute expression to decrease and the relative expression to diverge, while increasing promoter sensitivity (by decreasing K or n) had the opposite effect. Changes to promoter sensitivity were especially consequential for Msn2(T)*, which was effectively nonfunctional over a large range of promoter sensitivities where Msn2* and Msn2(A)* were still potent inducers (**Figure S12**). Except at low promoter sensitivities (high K or n) where expression differences between Msn2 mutants were always large, slow activation kinetics (low k_1_ or k_2_) or fast deactivation kinetics (high d_1_ or d_2_) caused the expression for Msn2(A)* and Msn2(T)* to diverge from the expression for Msn2*, while scaling the rate of transcription k_3_ had no effect on relative expression (**Figure 5B**). Decreasing the promoter transition rate (k_1_) caused the relative expression to diverge by increasing the relative contribution of TF affinity to the rate of transition from P_0_ to P_off_, while increasing d_1_ prevented any gains in P_off_ from accruing. Likewise, decreasing the on rate k_2_ or increasing the off rate d_2_ prevented gains in P_off_ from being converted to the productive state P_on_. These simulations show that it is possible for promoters with similar sensitivities but different kinetic parameters to have divergent responses to the Msn2 mutants, as was the case with the mid sensitivity promoters *CTT1* and *DCS2*.

We next used the gene expression model to explore how changes in nuclear localization dynamics may cause expression differences between the Msn2 mutants. We repeated the simulations described above, but for ideal pulses of each Msn2 mutant with varying amplitudes, durations, or interpulse durations (**Figure S13**). As with scaling the kinetic parameters, there was little scope to drive changes in the relative expression of low sensitivity promoters by changing the nuclear localization dynamics, though for higher sensitivity promoters, decreasing the amplitude of localization generally caused expression to diverge (**Figure 5C**), since decreasing the concentration of nuclear Msn2 increased the relative contribution of Msn2 affinity to the rate of transition from P_0_ to P_off_. Such behavior can be seen in our measurements of *RTN2*, where expression for the Msn2 mutants diverges with increasing pulse amplitudes, and *HSP12*, where expression converges (**Figure 4Ai**). In contrast, changing the duration of a single pulse of nuclear localization or the interpulse duration between multiple pulses had little effect on the relative expression of each hypothetical promoter (**Figure S13B – S13C**).

### Changing Msn2 affinity alters a cell’s ability to discriminate between stresses

Msn2(A)* induced low sensitivity promoters like *RTN2* and *TKL2* following pulsatile doses of nuclear localization where Msn2* and Msn2(T)* did not. We therefore predicted that it may similarly be better at inducing these genes in response to glucose starvation, which naturally causes sporadic pulses of Msn2 nuclear translocation (Hao and O’Shea, 2012). We therefore measured reporter expression for Msn2, Msn2(A), and Msn2(T)—all without CLASP or any mutations outside their DBDs—following glucose starvation and hyperosmotic shock, which causes an early, sustained pulse of Msn2 nuclear localization with a dose-dependent duration (**Figure 6 and Figure S14A**). Overall, expression of most promoters was highest for Msn2(A) and lowest for Msn2(T). While all three Msn2 mutants activated the low sensitivity genes *RTN2* and *TKL2* in response to hyperosmotic shock, only Msn2(A) activated them in response to glucose starvation. In essence, these genes lost their ability to discriminate between the stresses when the DNA binding affinity of Msn2 was increased. This was not due to differences in localization behavior, as both Msn2(A) and Msn2 localized similarly in response to either stress (**Figure S14B**). Excluding *DDR2*, which was most strongly induced by Msn2, there were few significant differences in the expression of the high sensitivity promoters between Msn2(A) and Msn2, which agrees with our model’s predictions that at a point there is little benefit to further increases in TF affinity.

**Figure 6.**
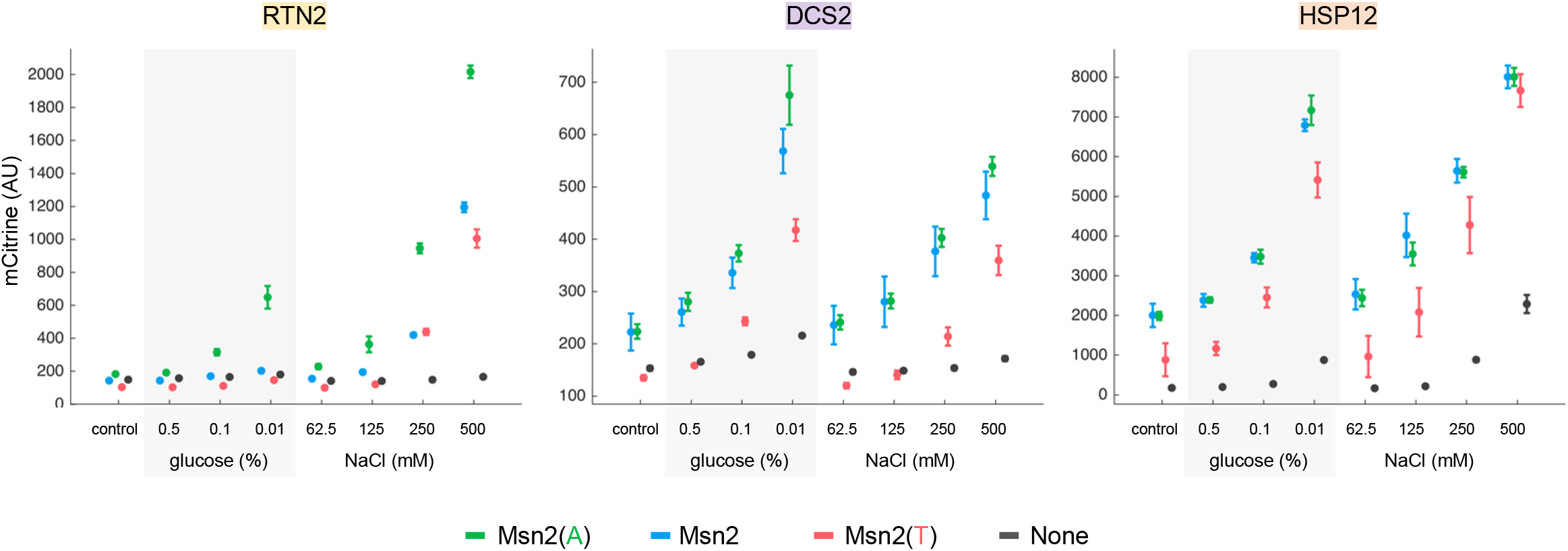
Changing Msn2 affinity can alter the ability of promoters to discriminate between stresses A.Fluorescent reporter expression following 2 hours of glucose starvation or hyperosmotic shock. All Msn2 mutants were expressed in the dCLASP system and had no mutations outside the DBD. Points and error bars represent the mean and standard deviation of fluorescence for at least three biological replicates. See also Figure **S14A**.

## DISCUSSION

Cells may encode environmental information in the temporal dynamics of TF activation and indeed previous work by Hansen and O’Shea showed that Msn2 target genes can decode the patterns of Msn2 nuclear localization generated by modulating PKA activity. However, other proteins downstream of PKA also regulate gene expression, including the TFs Hsf1, Sok2, and Dot6 and components of the mediator complex (Chang et al., 2004; Gutin et al., 2015; Lee et al., 2008; Pincus et al., 2014). Our measurements show that the signal decoding behavior of Msn2 target genes persists when Msn2* localization is controlled directly. The promoters act as filters of Msn2 localization dynamics: low sensitivity genes filtered out low amplitude, short duration, and pulsatile doses of nuclear Msn2, while high sensitivity genes are readily induced by nuclear Msn2, effectively integrating the nuclear Msn2 signal. Our results are broadly consistent with those of Hansen and O’Shea, though some promoters behaved differently than expected: *DCS2* and *RTN2* were less readily activated and *SIP18 D6*, which was previously reported to have a low amplitude threshold and high activation timescale, had intermediate values of both (**Figure 2C**). Accordingly, we observed no decoupling of amplitude threshold and activation timescale, which were linearly related for all promoters measured. Such differences may be due to differences in methodology. Beyond employing optogenetic control of Msn2 rather chemical control of PKA, we also used fluorescent reporters integrated at the URA3 locus rather than at the open reading frame (ORF) of each reporter gene.

Our light sweep experiments systematically probed how changes to the DNA binding affinity of Msn2 affected promoter decoding of its nuclear localization dynamics (**Figure 3C**). In general, increasing Msn2 affinity increased the expression of its target genes, making them more responsive to shorter, weaker, and pulsatile doses of nuclear Msn2, though these effects were much stronger for low sensitivity genes than high sensitivity genes. In fact, some low sensitivity genes lost the ability to discriminate between natural stresses when Msn2 affinity was increased (**Figure 6**), which is consistent with suggestions that the ability of TFs to bind DNA is tuned for function and high binding affinity is not necessarily optimal (Aditham et al., 2021; Crocker et al., 2016). A sensitivity analysis of our gene expression model indicated that—beyond affinity—slow activation kinetics or fast deactivation kinetics could cause a promoter to exhibit divergent responses to high and low affinity TF mutants. Moreover, a comparison of the *SIP18* mutants showed that, in many respects, the effect of increasing Msn2 affinity was similar to adding Msn2 binding sites to a promoter (**Figure S9**), though the relative ability to respond to pulsed versus continuous doses of nuclear Msn2 was set largely by the promoter. This is consistent with a proposal that *SIP18 A4* is more responsive to short or pulsed bursts of nuclear Msn2 than *SIP18 D6* because it facilitates Msn2 binding and subsequent chromatin remodeling at the TATA box (Hansen and O’Shea, 2015a).

The increase in gene expression typically associated with increasing Msn2 affinity was accompanied by a decrease in expression noise (**Figure S10**). Noisy expression can be beneficial as a source of phenotypic diversity between cells but limits the information transduction capacity of genes. Hansen and O’Shea previously identified a tradeoff between noise and control of gene expression: low sensitivity promoters filter out noisy TF activity, but respond to real signals with high levels of expression noise, while high sensitivity promoters have low expression noise, but are readily induced by noisy bursts of TF activity (Hansen and O’Shea, 2013, 2015b). Various strategies have been proposed for overcoming the effects of noise in decoding TF dynamics: integrating the response of multiple genes may allow cells to overcome the noisy expression of individual genes, recruiting chromatin regulators can fine-tune the information capacity of a given gene, and coordinated regulation by multiple TFs can provide finer control of gene expression (Benzinger et al., 2022; Gasch et al., 2017; Lin et al., 2015). Our results point to another potential mechanism for selectively activating genes with a single TF: concerted regulation of TF localization dynamics and DNA binding affinity, which could exploit promoter dependent differences in responsiveness to both modes of regulating TF activity. For example, our measurements of *RTN2* and *HSP12* (**Figure 4C**) show that a TF capable of flipping between high and low affinity DNA binding modes and exhibiting either pulsatile or continuous localization dynamics could tune the expression of low sensitivity genes, while maintaining the robust expression of high sensitivity genes (**Figure 7**). Such regulation may be beneficial in avoiding the activation of resource intensive or terminal cell fate genes when responding to mild stresses. Future studies are needed to determine if cells employ such strategies to coordinate stress-specific gene expression responses via TFs (like p53 and NF-κB) whose localization dynamics and DNA binding affinity are both subject to regulation.

**Figure 7.**
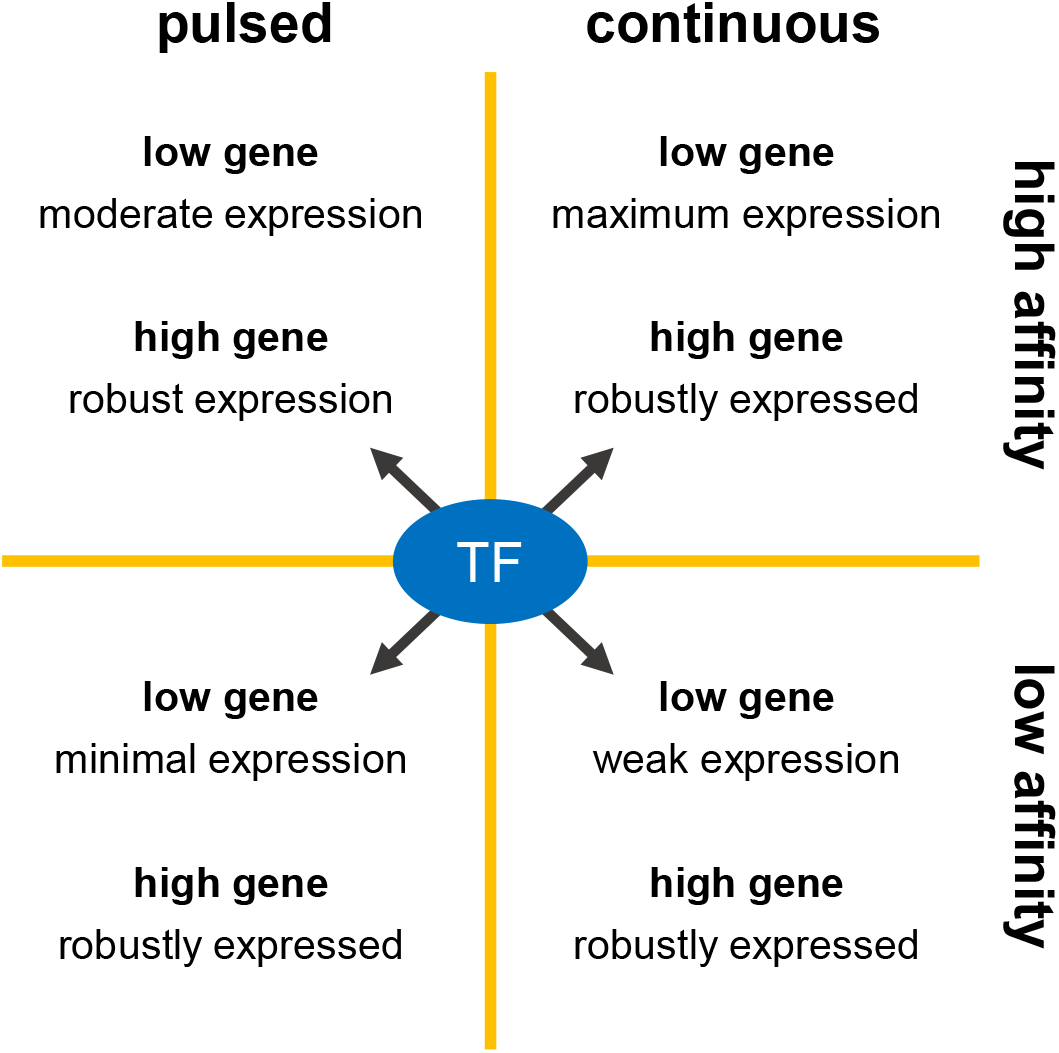
Concerted regulation of TF affinity and dynamics may facilitate improved control of gene expression A TF that could flip between high and low affinity binding modes and continuous and pulsatile nuclear localization dynamics could tune the expression of a low sensitivity gene over a wide range while maintaining the robust activation of a high sensitivity gene. Schematic is based on the single-cell expression measurements of **Figure 4C**, where a pulsed dose of the low affinity Msn2(T)* mutant minimally activated the low sensitivity gene RTN2, while a continuous dose of Msn2(T)* weakly activated RTN2. In contrast, a pulsed dose of the high affinity Msn2(A)* mutant moderately activated RTN2 and a continuous dose maximized RTN2 activation. Meanwhile, any of these doses of Msn2(T)* or Msn2(A)* were sufficient to robustly activate the high sensitivity gene *HSP12*.

Mechanistically, what drives the different behaviors of the high and low sensitivity promoters? Our gene expression model identified the affinity between a promoter and a TF as a key factor. Indeed, no low sensitivity promoter had more than two STREs within 500 bp of its ORF, while no high sensitivity promoter had fewer than four STREs in the same region (**Figure S5C**), which is consistent with observations that increasing the number of TF binding sites in a promoter generally increases its affinity for the TF and maximum expression (Lam et al., 2008; Sharon et al., 2012). Differences in nucleosome occupancy between the high and low sensitivity promoters may also drive differences in their behavior and manifest as differences in affinity by restricting access to STREs. Induction of the low sensitivity promoter *SIP18* involves a slow chromatin remodeling step before initiating transcription, as do mammalian genes that require sustained NF-κB activity (Hansen and O’Shea, 2013, 2015a; Sen et al., 2020). Likewise, Crz1 target promoters with low slope ratios are typified by a slow transition from an initial off state (P_0_) to an intermediate off state (P_off_) that is associated with high initial nucleosome occupancy at the promoter (Chen et al., 2020). Our low sensitivity reporters had both long activation timescales and low slope ratios (**Figure 2C and Figure 4B**) and were enriched for low predicted values of k_1_ and high predicted values of K^n^—all consistent with a slow transition step involving chromatin remodeling (**Figure S6B**). The general inability of Msn2(T)* to activate these promoters hints that it may be poor at initiating chromatin remodeling. Activation of latent enhancers in murine macrophages by NF-κB requires continuous NF-κB activity to disrupt histone-DNA interactions (Cheng et al., 2021) and competition with nucleosomes has also been implicated in the activation of yeast genes by Rap1 (Lickwar et al., 2012). In fact, the affinity of exposed Pho4 binding sites in the promoters of yeast phosphate response genes determines the level of phosphate starvation—and thus nuclear Pho4—needed to nucleate chromatin remodeling and activate gene expression (Lam et al., 2008). Further studies are needed to determine if the reduced DNA-binding ability of Msn2(T)* similarly limits its ability to compete with nucleosomes and initiate chromatin remodeling events.

Finally, we observed decay in our Msn2 nuclear localization time courses consistent with observations that nuclear accumulation of Msn2 triggers its degradation to overcome the growth defect caused by Msn2 hyperactivity (Durchschlag et al., 2004; Lallet et al., 2004; Sadeh et al., 2011). Msn2 degradation was greatest for long pulsatile light programs; the same may be true for natural stresses like glucose starvation that cause a drawn-out series of pulses of nuclear Msn2. Msn2 degradation caused the expression of low sensitivity genes to plateau for long duration light programs as nuclear Msn2 levels fell below the amplitude threshold needed for activation. Similar behavior in response to natural stress may cause transient activation of low sensitivity genes and sustained activation of high sensitivity genes. Our measurements revealed that Msn2 degradation began about 14 min after it first entered the nucleus—similar optogenetic approaches could prove useful in future efforts to study the dynamics of such processes.

## ACKNOWLEDGEMENTS

This work was supported by National Institutes of Health grant R35GM128873 and National Science Foundation grant 2045494 (awarded to M.N.M). Flow cytometry was enabled by the University of Wisconsin Carbone Cancer Center Support Grant P30 CA014520. Megan Nicole McClean, PhD holds a Career Award at the Scientific Interface from the Burroughs Wellcome Fund. Kieran Sweeney was supported by a NHGRI Training grant 5T32HG002760 to the Genomic Sciences Training Program at the University of Wisconsin-Madison. We thank Lindsey Osimiri and Hana El-Samad for providing us with CLASP and Edvard Grødem for building and modifying our optoPlates.

## AUTHOR CONTRIBUTIONS

K.S. and M.N.M. conceived of the study. K.S. collected and analyzed data. K.S. and M.N.M interpreted results and wrote the manuscript.

## METHODS

### Strain construction

The *Saccharomyces cerevisiae* strains used in this study were constructed from a base strain in the S288C background (MAT alpha his3D1 leu2D0 lys2D0 MET15 ura3D0). To identify the nucleus, the nuclear protein Nhp6a was tagged with the infrared fluorescent protein iRFP via URA3 pop-out. Briefly, this entailed tagging the C-terminal of Nhp6a using a caURA3 selective marker that was subsequently “popped out” by counterselection with 5-Fluoroorotic acid (5FOA) and a repair DNA template with homology to the genomic DNA regions immediately upstream and downstream of the caURA3 marker. To avoid interference with the Msn2-CLASP mutants, the native copy of Msn2 and its paralog Msn4 were deleted in the base strain, also by URA3 pop-out.

Reporters strains were then constructed from the base strain. Reporters were selected as follows: *HXK1, DCS2, SIP18, SIP18 A4, SIP18 D6, DDR2, TKL2, ALD3*, and *RTN2* were selected based on previous studies of the relationship between Msn2 nuclear localization and gene expression (Hansen and O’Shea, 2013, 2015a); *CTT1* was selected because it was used to study Msn2(S686A) activity (Reiter et al., 2013); and *HSP12* was selected because it was used to test Msn2-CLASP performance (Chen et al., 2020). A no reporter control strain featured GFP expressed under a bacterial promoter (*glpT*), which is silent in yeast. To create each reporter strain, the region 1000 bp upstream of the open reading frame of each reporter gene was amplified by PCR and inserted by Gibson assembly into an integrating plasmid such that it drove the expression of mCitrine. The reporter plasmids were screened by sequencing and integrated into the LEU2 locus of the base strain as previously described (Lee et al., 2015).

The Msn2 mutants were made using overlap PCR to mix and match, in a modular fashion, Msn2 domains with phosphomimetic mutations that were generated by PCR or purchased as gBlocks from IDT. The Msn2 mutants were then added by Gibson assembly to a CLASP plasmid without a cargo protein (pLO405), which was generously provided by Lindsey Osimiri and Hana El-Samad. Similarly, an equivalent dCLASP plasmid, lacking Zdk1 and yeLANS, was created for each Msn2 mutant by Gibson assembly. The Msn2-CLASP and Msn2-dCLASP plasmids were screened by sequencing and integrated into the URA3 locus of the reporter strains as previously described (Chen et al., 2020). The resulting transformants had inconsistent levels of mScarlet, suggesting that Msn2-CLASP sometimes integrated more than once, likely because regions of self-homology in the Msn2-CLASP plasmids were undergoing homologous recombination when transformed into yeast. Accordingly, we screened all transformants by flow cytometry for consistent, low mScarlet levels prior to use in the light sweep experiments.

Plasmid construction was done using DH5alpa competent cells. Yeast transformations were done using SC agar plates with appropriate auxotrophic or drug selection. After screening, yeast were frozen down and grown out on YPD plates for subsequent use in experiments. For all flow cytometry and microscopy experiments, yeast were grown in LFM (Sheff and Thorn, 2004). For practical reasons, strain construction was done under room lights, but all strains were incubated and stored in the dark.

### Blue light delivery

Blue light stimulation was done using an optoPlate (Bugaj et al., 2017) modified and calibrated as described previously (Grødem et al., 2020). Briefly, custom adaptors were designed in 3DS Max and 3D printed to mount the optoPlate upside-down over a 96 well plate on an inverted fluorescence microscope and the optoPlate software was modified to allow 1) communication with a microscope, 2) programming of the optoPlate with a plate map, an Excel spreadsheet recording the light pattern for each well, and 3) calibration of the LEDs. After calibration, the relationship between LED amplitude (0 – 255 AU) and irradiance was quantified (**FIGURE S2**).

### Light sweep experiments

#### Overview

In each light sweep experiment, reporter induction was measured for two Msn2-CLASP mutants, each subjected to 14 light programs with blue light pulses spanning a range of amplitudes, durations, and oscillatory patterns. As a control, reporter induction was measured for equivalent Msn2-dCLASP mutants subjected to 50 min of 100% light. As a batch control, *HXK1* expression was measured for Msn2*-CLASP subjected to both 50 min 100% light and no light. In total, this entailed imaging 32 wells per light sweep experiment. For a given reporter strain and pair of Msn2-CLASP mutants, three light sweep experiments were performed, one for each of three biological replicates.

#### Growth conditions

The cultures used for each light sweep experiment were grown over multiple days so that they reached mid-log phase by the morning of each experiment (day 0). On the evening of day -2, single colonies were picked into 100 uL LFM in 96 well plate and grown at 30 C overnight. On the evening of day -1, the resulting saturated cultures were serial diluted 1:7000 into 3 mL LFM and grown at 30 C for 15 hours. By morning of day 0, the diluted cultures reached mid-log phase and were used for one of three rounds of light sweep experiments throughout the day. Cultures for the first round of experiments were used immediately, while those for the second and third rounds were, respectively, diluted back 1:30 into 3 mL LFM and grown at 30 C for 3 – 4 hours or diluted 1:5 into 3 mL LFM and grown for 7 – 8 hours. All day 0 steps were done with 30 C LFM and in dark.

#### Microscopy

Light sweep experiments were done in optical 96 well plates (CellVis P96-1.5H-N) that were pretreated with concanavalin A (MP Biomedicals) to allow cells to adhere to the plate bottom and immersion oil to facilitate the use of an oil immersion microscope objective. Briefly, 30 uL of 2 mg/mL concanavalin A was added to each well being used, incubated at room temperature for 15 min, and removed, then the bottom of the 96 well plate was coated with immersion oi (Olympus, Type F). Cultures to be imaged were diluted to OD600 = 0.125 – 0.150 in LFM and plated in an optical 96 well plate which was then loaded onto the stage of an inverted fluorescence microscope (Nikon TiE) that was kept dark and at 30 C by an incubating enclosure. The cells were allowed to settle and adhere to the plate bottom for 15 min, at which point the media was removed, the cells were washed three times with LFM, and fresh LFM was added. The cells were then allowed to equilibrate for at least 10 min, during which time the optoPlate was mounted on top of the 96 well plate, connected to the microscope computer via USB, and configured to deliver the appropriate light dose to each well.

For each light sweep experiment, each culture was imaged every 2.5 min for 160 min using a Nikon TiE inverted microscope equipped with a 60x oil immersion objective, an automated stage, and CCD camera. The microscope was controlled by NIS-Elements and acquired 3 images for each of 32 wells per timepoint: an iRFP image of the nuclear marker (400 ms, Nikon Intensilight lamp with 540/45x 720/60m Cy5.5 filter cube, 1.5x gain, ND8, extended NIR mode), an mScarlet image of the Msn2-CLASP mutant (400 ms, Nikon Intensilight lamp with 560/40x 630/75m mCherry filter, ND8, 1.5x gain), and a YFP image of the mCitrine reporter (75 ms, Nikon Intensilight lamp with 510/20x 545/30m rsYFP (red-shifted YFP) filter cube, ND8, 1.5x gain). Focus was maintained using the Nikon Perfect Focus System (PFS). The rsYFP (red-shifted YFP) cube and 75 ms exposure at ND8 were selected to prevent light-induced localization of Msn2-CLASP when imaging. Occasionally, the PFS lost focus due to changes in height over the large plate area being imaged or the optoPlate shifting slightly; in these cases, focus was re-established using custom NIS-Elements scripts. When imaging commenced, NIS-Elements instructed the optoPlate to initiate, allowing the timelapse microscopy and light program to operate in sync. Each light program included a 10 min delay before the blue LEDs were activated such that basal fluorescence could be measured. This corresponded to the 0 min timepoint. The microscope instructed the optoPlate to turn off the LEDs in each well as it was imaged.

Over the course of the light sweep experiments, over 2300 .ND2 timelapse images were acquired, one for each of 32 wells per experiment. To automate the handling of this large amount of image data, the strain loaded in each well and the light program to which it was subjected were recorded in an spreadsheet (a “plate map”) that was saved with each set of images. Using custom MATLAB scripts, the plate map was subsequently used to automatically label the ND2 images and the single cell fluorescence measurements extracted from them.

#### Image analysis

Each light sweep experiment produced 32 .ND2 timelapse images with 65 frames and 3 channels per frame: iRFP, mScarlet, and mCitrine. Because the optoPlate was mounted on top of the optical 96 well plate, we did not acquire images with transmitted light such as phase contrast or DIC images from which to segment cells. Likewise, because the mCitrine and mScarlet signals varied substantially over the course of an experiment—mCitrine was induced and mScarlet moved in and out of the nucleus—we segmented the cells from the iRFP images of the nuclear marker. This was done using custom image processing code written in MATLAB. Images were loaded using the Bio-Formats MATLAB toolbox (Linkert et al., 2010), a Laplacian-of-Gaussian filter was applied to the iRFP images for blob enhancement, and a region of interest (ROI) representing each nucleus was segmented from the resulting high-contrast images using MATLAB’s circle finder. This process largely excluded cells whose nuclei were out of focus. The circles were then enlarged by a factor of two to define a region of interest representing each cell, while the cytoplasm was defined as the region within each cell but outside the nucleus. If highly overlapping cells were identified, the cell with the lower “metric” value from MATLAB’s circle finder was removed.

Using the ROIs defined during segmentation, the nuclear, cytoplasmic, and cellular fluorescence of each cell was quantified as the median pixel value of these regions in the raw iRFP, mScarlet, and mCitrine images. The background fluorescence in each channel was measured as the mode pixel value outside all cell ROIs. The resulting single cell measurements were labeled with strain and light program information from the plate map associated with each experiment, as well as time information extracted from the .ND2 metadata. Measurements associated with aberrant frames in each timelapse, for example, due to shutter timing mismatches or temporary loss of focus, were identified as outliers in the plot the median iRFP of all cells versus frame and removed. Fluorescence differences due to long-term fluctuations in the Intensilight lamp intensity were corrected based on weekly lamp irradiance measurements. Photobleach correction was applied to the mCitrine measurements, but not mScarlet as the dynamic Msn2 localization timecourses were not amenable to this approach. The median background mScarlet and mCitrine level per experiment was subtracted from the single cell mScarlet and mCitrine measurements, respectively. Msn2 nuclear localization was quantified as nuclear mScarlet divided by cytoplasmic mScarlet. Basal Msn2 localization, quantified as the median Msn2 localization before the 0 min timepoint, was subtracted from the single cell Msn2 nuclear localization measurements. The Msn2 localization measurements were then normalized to the maximum observed level of nuclear localization.

The Msn2 localization and mCitrine induction timecourses for each combination of Msn2-CLASP mutant and reporter were captured for three biological replicates across three separate experiments. As a result, the precise time at which each frame was captured varied by experiment. This was exacerbated in cases where refocusing was needed and led to issues when averaging fluorescence measurements across the three experiments, especially when a frame for one of the replicates was dropped. To account for these issues, the population level Msn2-CLASP localization and mCitrine induction timecourses per condition were calculated by 1) taking the median localization or fluorescence measurement of all cells per frame, 2) assigning these values to bins representing 2.5 min segments of time, 3) filling in any missing values by linear interpolation. The mCitrine measurements for each replicate were smoothed using a 5-point moving average filter. Because Msn2(A)*-CLASP, Msn2*-CLASP, and Msn2(T)*-CLASP localize similarly in response to light, the area under curve of Msn2 localization per condition was calculated from the mean localization of all three Msn2 mutants across all experiments. Measurements were plotted using the Gramm data visualization toolbox (Morel, 2018).

#### Flow cytometry experiments with natural stress

As with the light sweep experiments, stains used for flow cytometry experiments were grown over the course of three days. On the evening of day -2, four colonies of each strain were picked into 100 uL of LFM in a 96 well plate and incubated at 30 C overnight. On the evening of day -1, the resulting saturated cultures were diluted 1:1600 into 200 uL LFM in a new 96 well plate and incubated at 30 C for 14 hours. On the morning of day 0, 20 uL of each culture was aliquoted into 8 plates containing 140 uL 30 C LFM and incubated 4 hours at 30 C. The cultures were pelleted by centrifuging for 5 min at 3200 rpm and then forcefully tipping out the supernatant. The pellets were resuspended in control media (30 C LFM), hyperosmotic shock media (30 C LFM with 0.5, 0,25,0.125, or 0.0625 M NaCl), or glucose deficient media (30 C LFM with 0.5, 0.1, or 0.01% glucose) and grown for 2 hours at 30 C. To arrest translation, 40 uL of 0.5 mg/mL cycloheximide was added to each culture and allowed to incubate at 30 C for 30 min, at which point 20 uL of each arrested culture was added to 140 uL 4 C PBS 0.1% tween in a 384 well plate.

The cells were then measured with an Attune NxT flow cytometer equipped with an autosampler: mScarlet was measured with 561 nm excitation light and a 585/16 nm filter and mCitrine was measured with a 488 nm excitation light and 590/40 nm filter. Rainbow beads were used to ensure day-to-day consistency in intensity measurements. The flow cytometry measurements were processed and analyzed using custom MATLAB scripts. Measurements were imported from FCS files using the fca_fcsread loader (Balkay, 2022) and automatically labeled from a plate map, an Excel spreadsheet file containing the strain and condition information for each well of the 384 well plate. The measurements were gated to remove debris and doublets and the median fluorescence intensity (MFI) was calculated for mScarlet and mCitrine per well. Measurement were plotted using the Gramm data visualization toolbox (Morel, 2018).

#### Gene expression model

The gene expression model (**Figure 2D**) features three promoter states and represents the production of mature YFP (mYFP) as a function of Msn2 with the following ordinary differential equations:

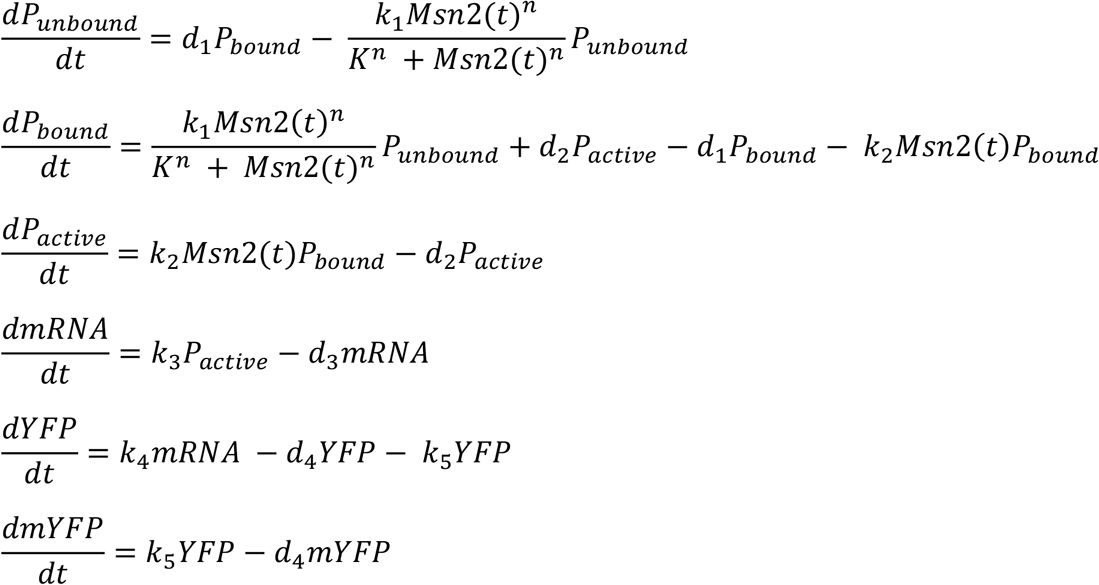

This three-state model of promoter activation was chosen based on previous reports that a transition between two off states (P_0_ and P_off_) captures the behavior of slow promoters like *SIP18* by representing chromatin remodeling steps needed for activation (Chen et al., 2020; Hansen and O’Shea, 2013; Sen et al., 2020). To capture the switch-like activation of some promoters, we modeled this transition with a Hill function, where K and n capture the half-maximum point and slope of the curve relating nuclear Msn2 concentration to the rate of promoter transition. K^n^ is related to the binding affinity between the promoter and Msn2, which is determined by the sequence, number, and location of Msn2 binding sites in the promoter as well as other factors like competition and nucleosome occupancy (Hansen and O’Shea, 2015a; Lam et al., 2008; Sharon et al., 2012; Stewart-Ornstein et al., 2013).

To parameterize the models, pooled Msn2 localization measurements (**Figure 2B**) were interpolated and used as the input Msn2(t) and the predicted YFP level (mYFP) was fit to the time-resolved localization measurements for reporter and Msn2 DBD mutant across all light programs. More specifically, we calculated expression for each light program for 100,000 parameter sets obtained by Latin hypercube sampling in which the promoter-specific parameters were allowed to vary over the following ranges: k_1_, d_1_, k_2_, and k_3_ from 10^−3^ – 10^2^, K from 1 – 10^4^, n from 0.5 – 4, and d_2_ from 10^−4^ – 10^2^. Global parameter values d_4_ = 0.08, k_4_ = 15, d_4_ = 0.001, and k_5_ = 0.06 were taken from the literature (Hansen and O’Shea, 2013). We then ranked the parameter sets by how well they minimized the residual sum of squares error between the predicted and measured expression across all light programs.

