## Supplemental Material for "Transcription Factor Localization Dynamics and DNA Binding Drive Distinct Promoter Interpretations"

### SUPPLEMENTARY FIGURE CAPTIONS

Figure S1. Screening Msn2 mutants.

A. Msn2 mutants were screened by their ability to localize in response to light. Using an optoPlate mounted on an inverted fluorescence microscope, cells were exposed to at least 5 min of 128 AU blue light and imaged. Bars and error bars represent mean and standard deviation nuclear/cytoplasmic Msn2 for two biological replicates. Msn2 mutants are named following the convention Msn2(NES|NLS|DBD) in which the residues highlighted in **Figure 1C** are mutated to either alanine (A) or glutamic acid (E) or left as WT. Mutations are color-coded red if they inhibit their respective domains. For example, Msn2(A|4E|WT) denotes a mutant with an inhibitory S288A mutation in its NES, inhibitory S582E, S620E, S625E, and S633E mutations (4E) in the NLS and a WT DBD. Msn2(WT|4E|WT) was selected from the screen and renamed Msn2* for convenience.
B. Screening Msn2-CLASP mutants by their ability to induce fluorescent reporters expressed under Msn2 target promoters. Reporter strains with Msn2-CLASP or Msn2-dCLASP mutants were exposed to 2 hours of continuous blue light (255 AU) and reporter fluorescence was measured by flow cytometry. Fold-change reporter induction was calculated relative to a dark treatment of each replicate. Bars and error bars represent the mean and standard deviation of 4 biological replicates.
C. The relationship between optoPlate input value (0 – 255 AU) and light output was measured as described previously (Grødem et al., 2020).
D. Fluorescence microscopy images showing localization of Msn2 and Msn2* with both CLASP and dCLASP in the dark and in response to 128 AU blue light with 2 s on 1 s off pulse-width modulation (PWM).
E. Histograms show nuclear Msn2 levels prior to illumination across at least 36 experiments for constructs with both CLASP and dCLASP: long right tail for Msn2 constructs reflects stochastic pulsing into the nucleus due to active NLS of Msn2. In contrast, Msn2* constructs have an inactivated NLS and undergo less stochastic pulsing into the nucleus. Overall, Msn2 constructs with CLASP are brighter than those with dCLASP, likely because cytoplasmic sequestration of Msn2 by CLASP prevents degradation triggered by localization to nucleus (see **Figure 2E**).

Figure S2. Comparison of expression measurements and model fits for low sensitivity promoters

Comparison of expression measurements and model fits for *RTN2*, *TKL2*, *ALD3*, and *SIP18*. Light solid lines and shaded regions show the mean and standard deviation of mCitrine fluorescence (AU) for at least three biological replicates. Dark dashed lines and shaded regions show the mean and 95% confidence interval of predicted mCitrine expression for the top 10 parameter sets for each reporter and Msn2 DBD mutant.

Figure S3. Comparison of expression measurements and model fits for mid sensitivity promoters

Comparison of expression measurements and model fits for *CTT1*, *SIP18 D6*, and *DCS2*. Light solid lines and shaded regions show the mean and standard deviation of mCitrine fluorescence (AU) for at least three biological replicates. Dark dashed lines and shaded regions show the mean and 95% confidence interval of predicted mCitrine expression for the top 10 parameter sets for each reporter and Msn2 DBD mutant.

Figure S4. Comparison of expression measurements and model fits for high sensitivity promoters

Comparison of expression measurements and model fits for *SIP18 A4*, *DDR2*, *HXK1*, and *HSP12*. Light solid lines and shaded regions show the mean and standard deviation of mCitrine fluorescence (AU) for at least three biological replicates. Dark dashed lines and shaded regions show the mean and 95% confidence interval of predicted mCitrine expression for the top 10 parameter sets for each reporter and Msn2 DBD mutant.

Figure S5. Light sweep experiment details

A. Each light sweep experiment involved 16 light programs. Light programs 1 – 5 featured 50 min pulses of blue light with amplitudes of 0, 20, 45, 85, and 128 AU. Light programs 6 – 9 featured pulses of 128 AU blue light with durations of 10, 20, 30, and 40 min. Light programs 10 – 12 featured 2, 6, and 10 five min pulses of 128 AU blue light with interpulse durations of 5 min. Light programs 13 and 14 featured 6 five min pulses of 128 AU blue light separated by 10 and 15 min, respectively. Light programs 15 and 16 (not shown) feature full-light doses (same as light program 5) for a dCLASP control with the appropriate Msn2 mutant (see **Figure S5D**) and a batch control, respectively. All light programs included 2s off, 1 s on pulse width modulation of the light dose.
B. Localization timecourses for all three Msn2 DBD mutants (with CLASP) in response to the above light programs. Lines and shaded regions represent the mean and standard deviation of Msn2 localization from at least 36 light sweep experiments per mutant.
C. Schematic depictions of Msn2 target promoters showing the location of the TATA box and each Msn2 binding site (STRE), which has the sequence AGGGG.
D. Maximum promoter expression in full light for each Msn2 DBD mutant with either CLASP (light program 5) or dCLASP (light program 15) show that light-induced gene expression is due to optogenetic control of each Msn2 DBD mutant via CLASP. Bars and error bars represent the mean and standard deviation of expression for at least three biological replicates.

Figure S6. Gene expression model details and calculating amplitude thresholds and activation timescales

A. $K^n$ captures both the half-maximum point and slope of the curve relating nuclear Msn2 concentration to the rate at which the promoter transitions from $P_{off}$ to $P_{on}$. He we depict of how individually scaling the K and n terms in the gene expression model affects the rate of promoter transition (from $P_0$ to $P_{off}$), which we take to represent chromatin remodeling at the promoter.
B. Predicted promoter parameters calculated by fitting the light sweep experiment measurements for Msn2* to the gene expression model. Fit results are shown in **Figures S2 – S4**. Points and error bars show the mean and 95% confidence interval of each parameter for the top 0.1% of parameter sets (top 100) for each promoter.
C. Schematic depicting the calculation of amplitude thresholds and activation timescales for each Msn2 DBD. We used the gene expression model to calculate the maximum level of $k_3P_{on}$ achieved following ideal pulses of nuclear Msn2* with a fixed amplitude (100%) and varying durations (0 – 50 min) or a fixed duration (50 min) and varying amplitudes (0 – 100%). We then calculated an amplitude threshold and activation timescale for Msn2* (dashed blue lines), which respectively describe the amplitude and duration of nuclear Msn2* needed to reach the half the maximum level of $k_3P_{on}$ attained in response to a 50 min 100% amplitude pulse of nuclear Msn2* (dotted blue lines). We next calculated amplitude thresholds and activation timescales for Msn2(A)* and Msn2(T)*, which respectively describe the amplitude and duration of nuclear Msn2(A*) (dashed green line) and nuclear Msn2(T)* (dashed red line) needed to achieve the half-maximum level of $k_3P_{on}$ attained by Msn2* (dotted blue line). Solid lines and shaded regions represent the mean and 95% confidence interval of predicted $k_3P_{on}$ for the top 10 (0.01%) parameter sets of *SIP18 A4*. We calculated these thresholds based on $k_3P_{on}$ rather than $P_{on}$ because $P_{on}$ varied substantially between parameter sets, as the rate of mRNA production ($k_3$) was allowed to vary widely when parameterizing the model. Scaling $P_{on}$ by $k_3$ produced a measure of promoter activation ($k_3P_{on}$) with drastically reduced variation. The model overestimated expression for short pulses of Msn2(A)* for *RTN2* and *TKL2*. Accordingly, levels of $k_3P_{on}$ for *RTN2* and *TKL2* were forced to zero for pulses of Msn2(A)* less than or equal to 10 min when calculating their relative activation timescales.

Figure S7. Quantifying localization triggered degradation of Msn2

A. Light sweep experiment in which every frame was taken at a different position indicates that decay in Msn2 localization time-coureses is not due to photobleaching. Lines and shaded regions represent mean and standard deviation of pooled localization measurements for three biological replicates each of Msn2*-CLASP and Msn2(A)*-CLASP. Experiments were performed in a no reporter control strain (*glpT*) to avoid mCitrine crosstalk into mScarlet channel.
B. Light sweep experiment for Msn2(D)*-CLASP, which features an S686D mutation in Msn2 that inhibits DNA-binding (Reiter et al., 2013). Lines and shaded regions represent mean and standard deviation of Msn2(D)* localization from 12 experiments representing 3 biological replicates each for 4 reporter strains. Msn2* localization acquired in separate experiments (see **Figure S5B**) is plotted for comparison.
C. The difference between the measured and ideal Msn2* localization timecourses (**Figure 2E**) shows that localization triggered degradation of Msn2* begins about 14 min after it first enters the nucleus (at t = 0 min). Peaks before 14 min are due to slight (single frame) timing mismatches between the measured and ideal timecourses; larger peaks of this nature were manually excluded from the plot for clarity.
D. Msn2 localization measurements for the first and second pulses of light programs 11, 13, and 14 indicate that Msn2 degradation was triggered by nuclear localization but continued outside the nucleus. Light programs 11, 13, and 14 each feature six 5 min pulses of nuclear Msn2 separated by interpulse durations of 5, 10, and 15 min, respectively. The first pulses of these light programs were concurrent and had no significant differences in amplitude, but the second pulses were staggered and their amplitudes decreased as the time since the first pulse increased. Boxplots show localization measurements for each pulse pooled from 79 separate measurements of Msn2(A)*, Msn2*, and Msn2(T)*. Significance was calculated by performing two-sample t-tests between each pair of pulses.

Figure S8. Msn2 DBD mutants details

A. Schematic depiction of the DNA binding domains (residues 645 – 704) of Msn2*, Msn2(A)* and Msn2(T)*. Canonical recognition residues that contact DNA are highlighted in grey. Msn2* has a WT DBD, Msn2(A)* features an S686A mutation within the second zinc finger, and Msn2(T)* features three mutations (S671T, N672G, R674K) within the linker region of the Msn2 DBD.
B. Amplitude thresholds and activation timescales were calculated as described previously (see **Figure S6C**). Bars and error bars show the mean and 95% confidence interval of amplitude and duration threshold for the top 10 parameter sets for each promoter and Msn2 DBD mutant. No thresholds are reported for Msn2(T)* where it unable to activate a given promoter in most cells.
C. Comparison of measured and predicted expression of select high and low sensitivity promoters. Points and error bars show mean and standard deviation of measured expression for three biological replicates. Solid lines and shaded regions show the mean and 95% confidence interval of predicted expression for the measured pulses of Msn2 nuclear localization. Dashed lines and shaded regions show the mean and 95% confidence interval of predicted expression for 100% amplitude ideal pulses of each Msn2 DBD mutant with durations ranging from 0 – 50 min. In both cases, simulations were performed using the top 10 parameter sets for each reporter and Msn2 DBD mutant. Predicted expression of the low sensitivity promoters plateaued when the measured Msn2 localization timecourses were used as the model input, but continued increasing with pulse duration when the ideal Msn2 localization timecourses were used—suggesting that degradation of Msn2 beyond 30 min caused the plateau and thus acted as a timer on expression.

Figure S9. Comparison of *SIP18* mutants

Expression of *SIP18*-based promoters following a 30 min 100% amplitude pulse of nuclear Msn2 (continuous) or six 5 min 100% amplitude pulses with 5 min interpulse durations (pulsed). Lines and shaded regions represent the mean and standard deviation of expression for three biological replicates. Schematic depictions of each promoter (top) show TATA box and STREs located within region 750 bp upstream of mCitrine. *SIP18 A4* was created from *SIP18* by replacing its native STREs with a cluster of four STREs proximal to the TATA box. Similarly, *SIP18 D6* was created by replacing the native STREs of *SIP18* with a cluster of six STREs distal to the TATA box.

Figure S10. Decreased Msn2 affinity and pulsatile nuclear localization dynamics increase gene expression noise

A. Noise levels following a 30 min 100% amplitude pulse of nuclear Msn2 (circles) or six 5 min 100% amplitude pulses with 5 min interpulse durations (triangles). Noise level was calculated per timepoint as $\sigma^2/\mu^2$ of all single-cell mCitrine measurements. Each point represents the average noise level of a single biological replicate for all timepoints greater than 120 min.
B. Noise ($\sigma^2/\mu^2$) versus maximum expression (AU) following a 50 min 100% amplitude pulse of each Msn2 DBD mutant. Each point represents the mean of three biological replicates. Noise was calculated for each replicate as the mean noise level for times greater than 120 min.

Figure S11. Incorporating differences between Msn2 DBD mutants into gene expression model

A. Calculating scaling factors α and β to represent differences in binding affinity between the Msn2 DBD mutants. We repeatedly simulated gene expression using the top 10 parameter sets for each reporter and Msn2(A)* while scaling α and β to minimize the error between the predicted expression and the measured expression for Msn2* or Msn2(T)* across all reporters and light programs. Error was quantified for each promoter, parameter set, and pair of α and β values as the residual sum of squares error across all light programs (RSS) divided by the overall minimum RSS of each promoter ($RSS_{min}$). Lines show mean of normalized error ($RSS/RSS_{min}$) across all promoters versus α for each value of β (different colors). Based on the calculations we chose scaling values of α =1 and β = 1 for Msn2(A)*, α =1.75 and β = 1 for Msn2*, and α =1 and β = 1 for Msn2(T)* to represent differences in binding affinity between the Msn2 DBD mutants.
B. Comparison of measured and predicted expression using the top 10 parameter sets for Msn2(A)* for each promoter and the chosen values of α and β. Points and error bars represent mean and standard deviation of measured expression for three biological replicates. Lines and shaded regions represent the mean and 95% confidence interval of predicted expression.

Figure S12. Predicted expression of selected hypothetical promoters for each Msn2 DBD mutant

Maximum predicted expression of hypothetical promoters in response to a 50 min 100% amplitude ideal pulse of nuclear Msn2(A)*, Msn2*, or Msn2(T)*. At a given value of K or n, each kinetic parameter ($k_1$, $d_1$, $k_2$, $d_2$, or $k_3$) was individually varied from $4^{-4}$ – $4^4$ while all other kinetic parameters were fixed at a value of one. K was allowed to vary from $2^0$ – $2^8$ and n was set to 0.5, 1, 1.5, 2, or 4. All values were predicted using the three-state promoter model depicted in **Figure 5A** and the values α = 1 and β = 1 for Msn2(A)*, α = 1.75 and β = 1 for Msn2*, α = 1.75 and β = 1.75 for Msn2(T)*.

Figure S13. Predicted expression of hypothetical promoters versus nuclear localization

Predicted expression of selected hypothetical promoters in response to (A) a 50 min pulse of each Msn2 DBD mutant with amplitudes ranging from 0 – 100% (B) 100% amplitude pulses of each Msn2 DBD mutant with durations ranging from 0 – 50 min, and (C) six 5 min 100% amplitude pulses of each Msn2 DBD mutant with interpulse durations ranging from 0 – 15 min. Top set of plots shows maximum expression of each hypothetical promoter, while bottom set of plots shows maximum promoter expression for each Msn2 DBD mutant normalized to maximum for Msn2*. In each 3 x 3 grid of plots, promoter affinity and given kinetic parameter were allowed to vary while all other kinetic parameters were fixed at one. For all plots, $n = 1$. All values were predicted using the three-state promoter model depicted in **Figure 5A** and the values $\alpha = 1$ and $\beta = 1$ for Msn2(A)*, $\alpha = 1.75$ and $\beta = 1$ for Msn2*, $\alpha = 1.75$ and $\beta = 1.75$ for Msn2(T)*.

Figure S14. Reporter expression and Msn2 localization following glucose starvation and hyperosmotic shock.

A. Fluorescent reporter expression following 2 hours of glucose starvation or hyperosmotic shock. All Msn2 mutants were expressed in the dCLASP system and had no mutations outside the DBD. Points and error bars represent the mean and standard deviation of fluorescence for at least three biological replicates.
B. Fluorescence microscopy measurements of nuclear/cytoplasmic localization for Msn2 and Msn2(A) following glucose starvation or hyperosmotic shock. Each violin plot represents single-cell localization measurements over a 12.5 min time window for three biological replicates of each Msn2 mutant. Msn2 mutants used here were in the dCLASP system and had no mutations outside the DBD.

### Figure S1

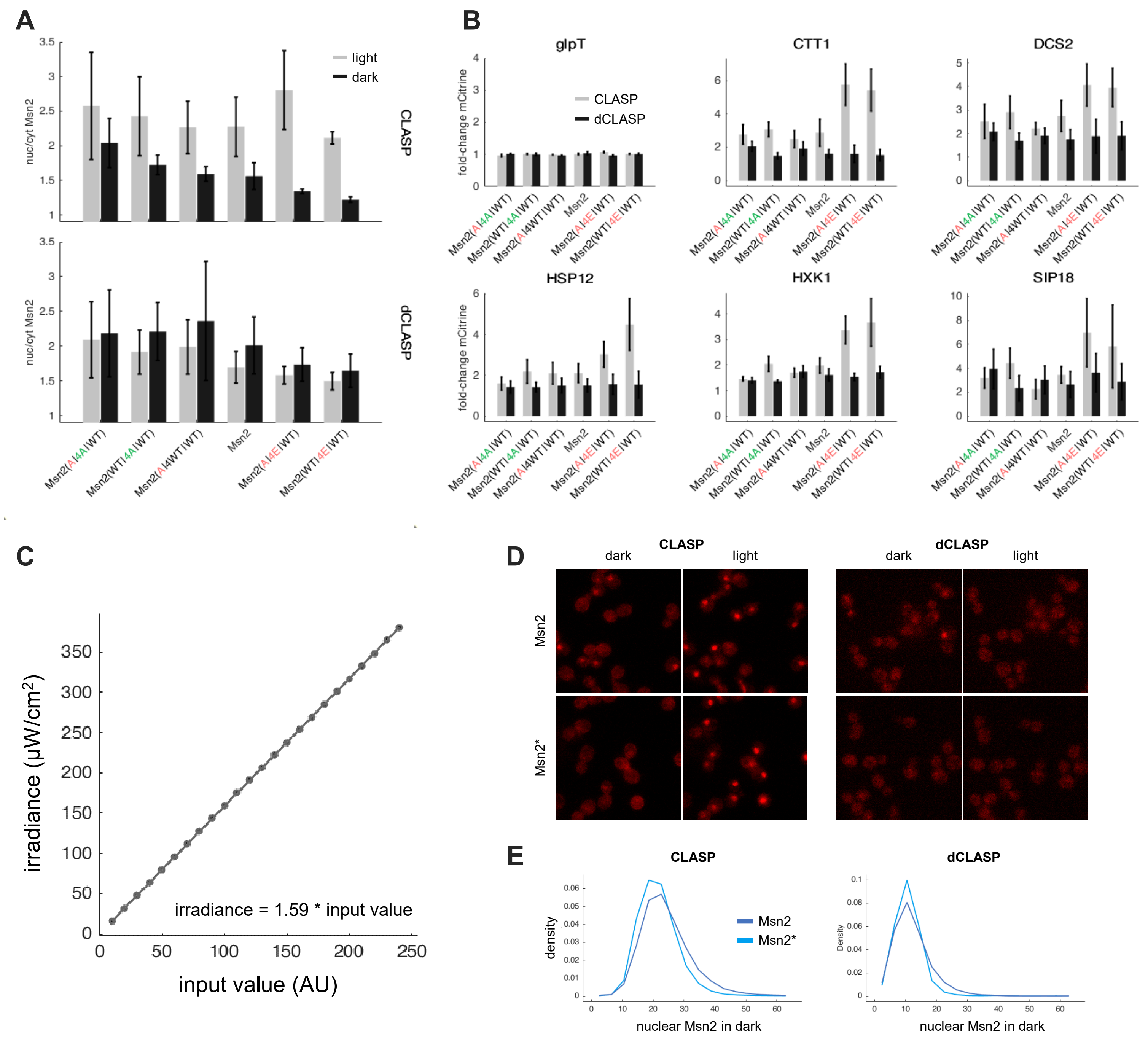
A
light
dark
nuc/cyt Msn2
CLASP
dCLASP
Msn2(A|4A|WT)
Msn2(WT|4A|WT)
Msn2(A|4WT|WT)
Msn2
Msn2(A|4E|WT)
Msn2(WT|4E|WT)
B
glpT
CTT1
DCS2
HSP12
HXK1
SIP18
fold-change mCitrine
CLASP
dCLASP
C
irradiance (μW/cm2)
input value (AU)
irradiance = 1.59 * input value
D
CLASP
dCLASP
dark
light
Msn2
Msn2*
E
CLASP
dCLASP
density
Density
Msn2
Msn2*
nuclear Msn2 in dark

### Figure S2

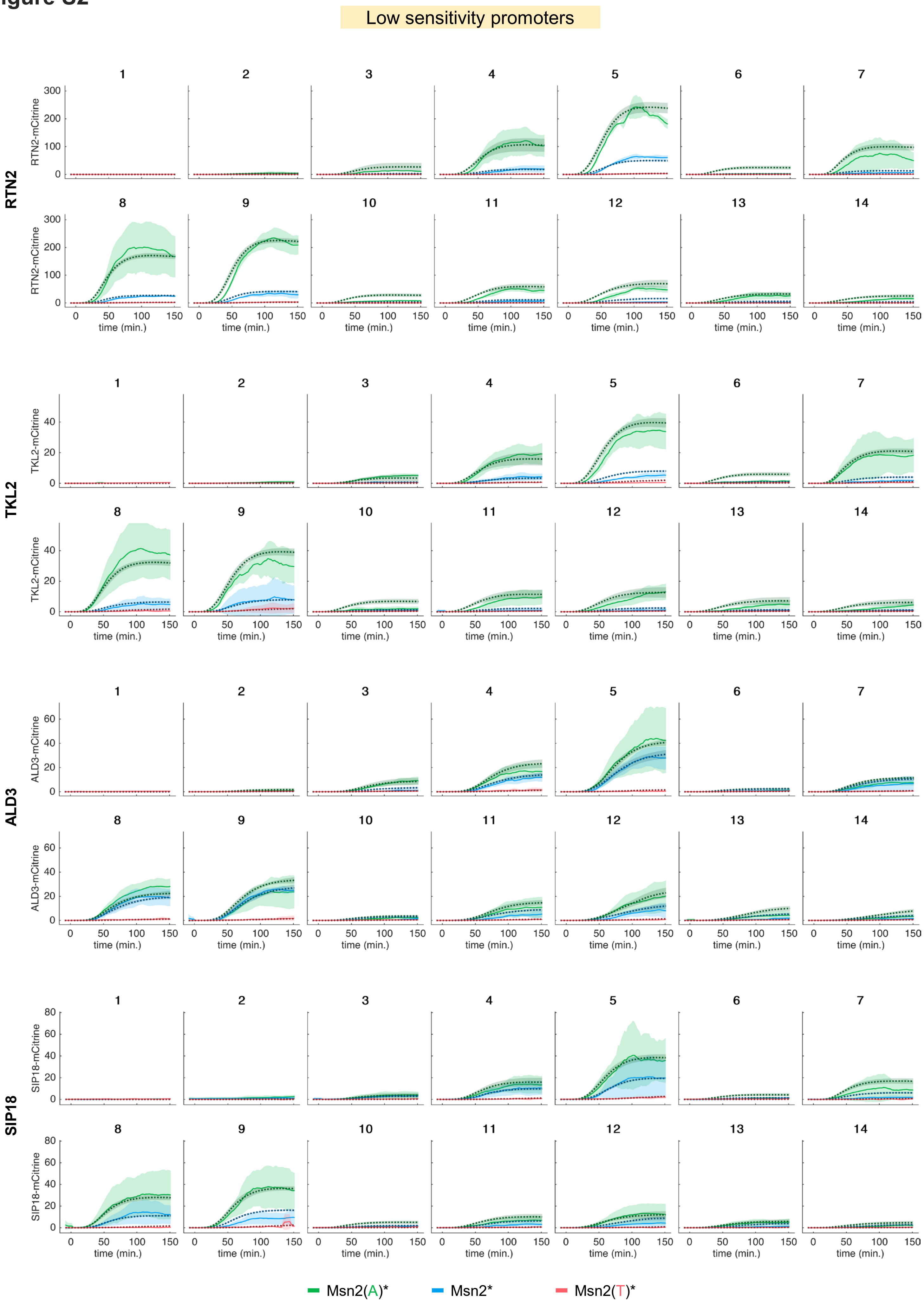

Low sensitivity promoters
RTN2
TKL2
ALD3
SIP18
RTN2-mCitrine
TKL2-mCitrine
ALD3-mCitrine
SIP18-mCitrine
time (min.)
Msn2(A)*
Msn2*
Msn2(T)*

#### Figure S3

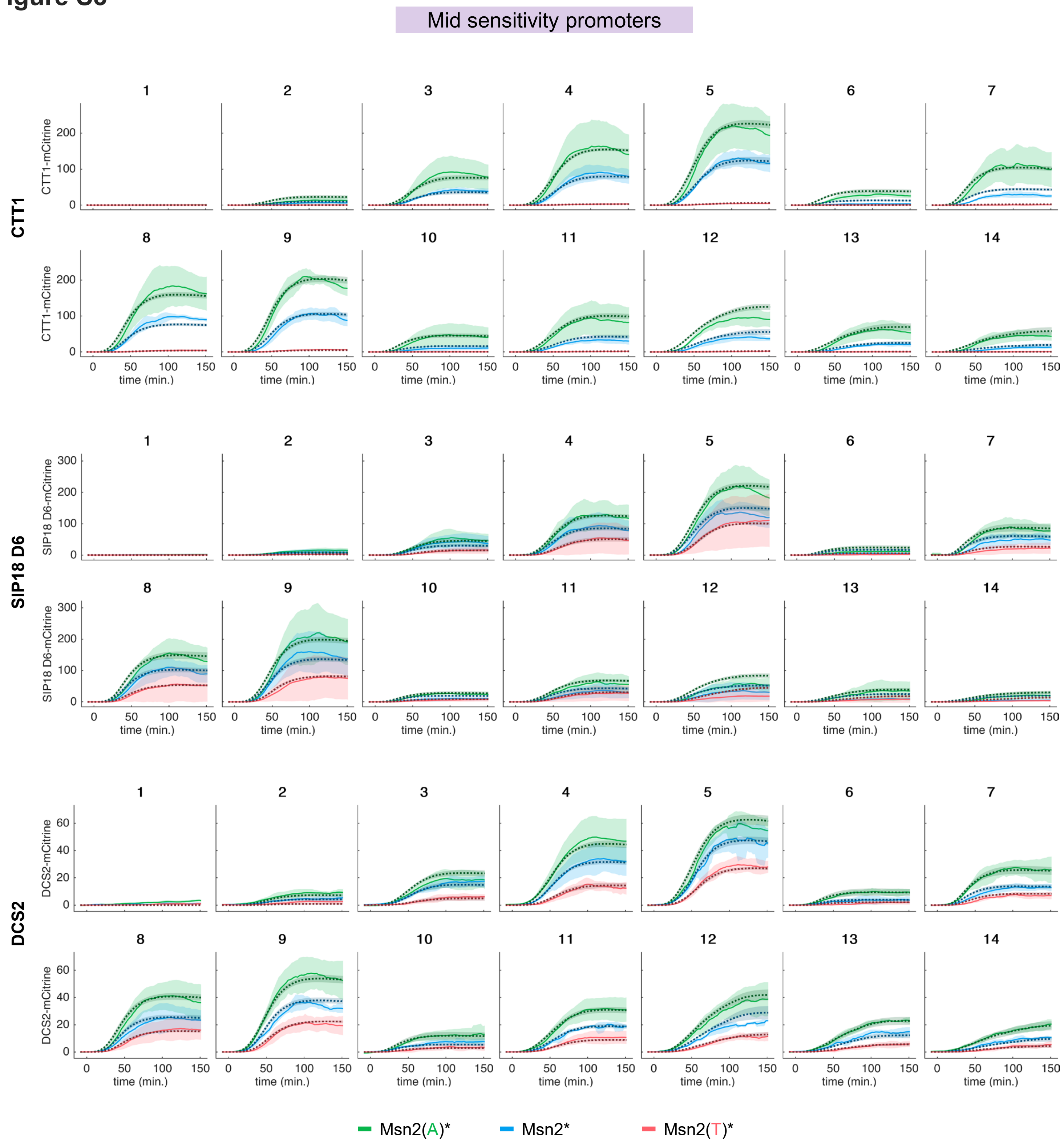

Mid sensitivity promoters
CTT1
CTT1-mCitrine
SIP18 D6
SIP18 D6-mCitrine
DCS2
DCS2-mCitrine
1
2
3
4
5
6
7
8
9
10
11
12
13
14
0
50
100
150
200
300
20
40
60
time (min.)
Msn2(A)*
Msn2*
Msn2(T)*

### Figure S4

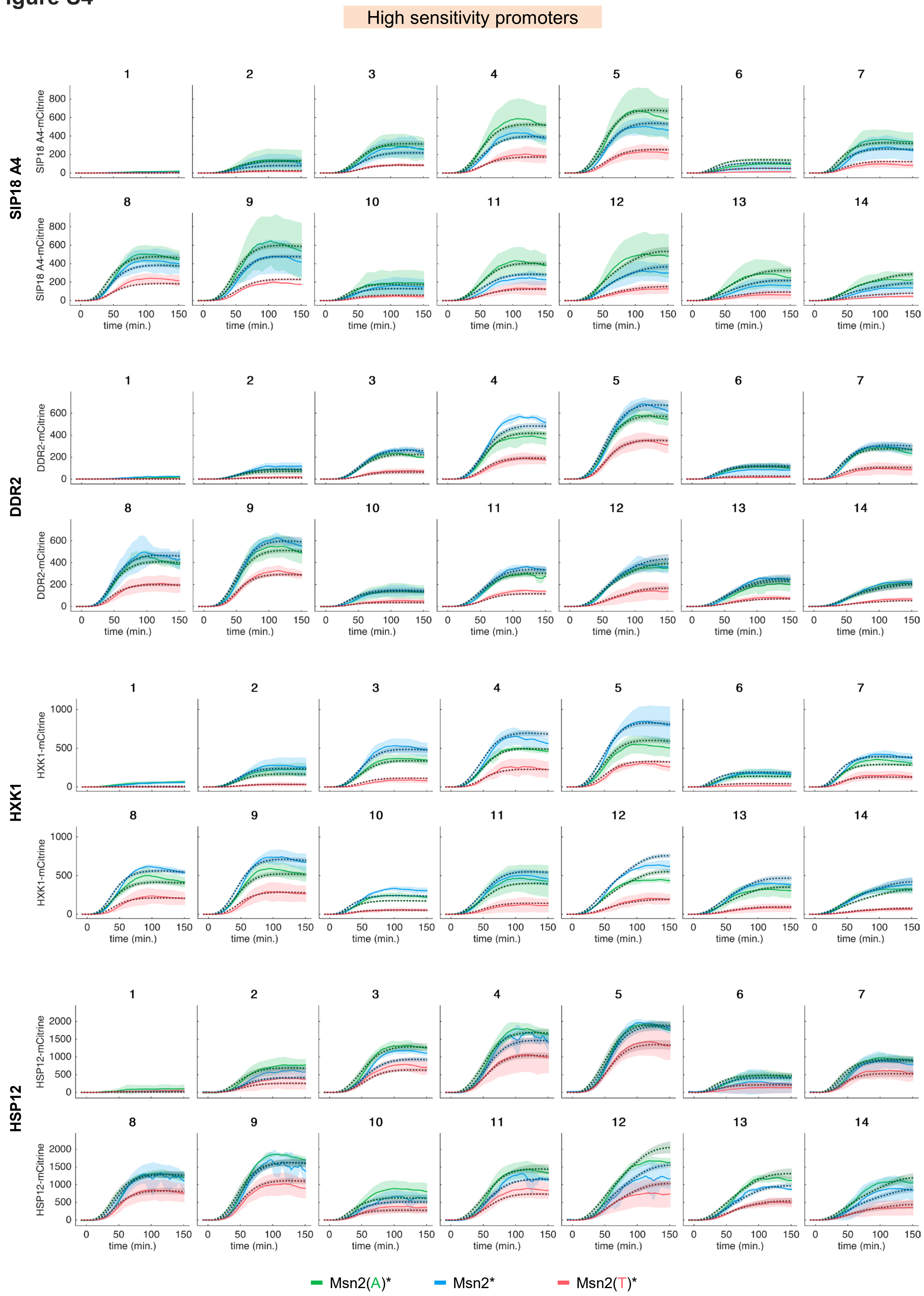
High sensitivity promoters
SIP18 A4
SIP18 A4-mCitrine
DDR2
DDR2-mCitrine
HXK1
HXK1-mCitrine
HSP12
HSP12-mCitrine
1
2
3
4
5
6
7
8
9
10
11
12
13
14
time (min.)
Msn2(A)*
Msn2*
Msn2(T)*

**Figure S5**

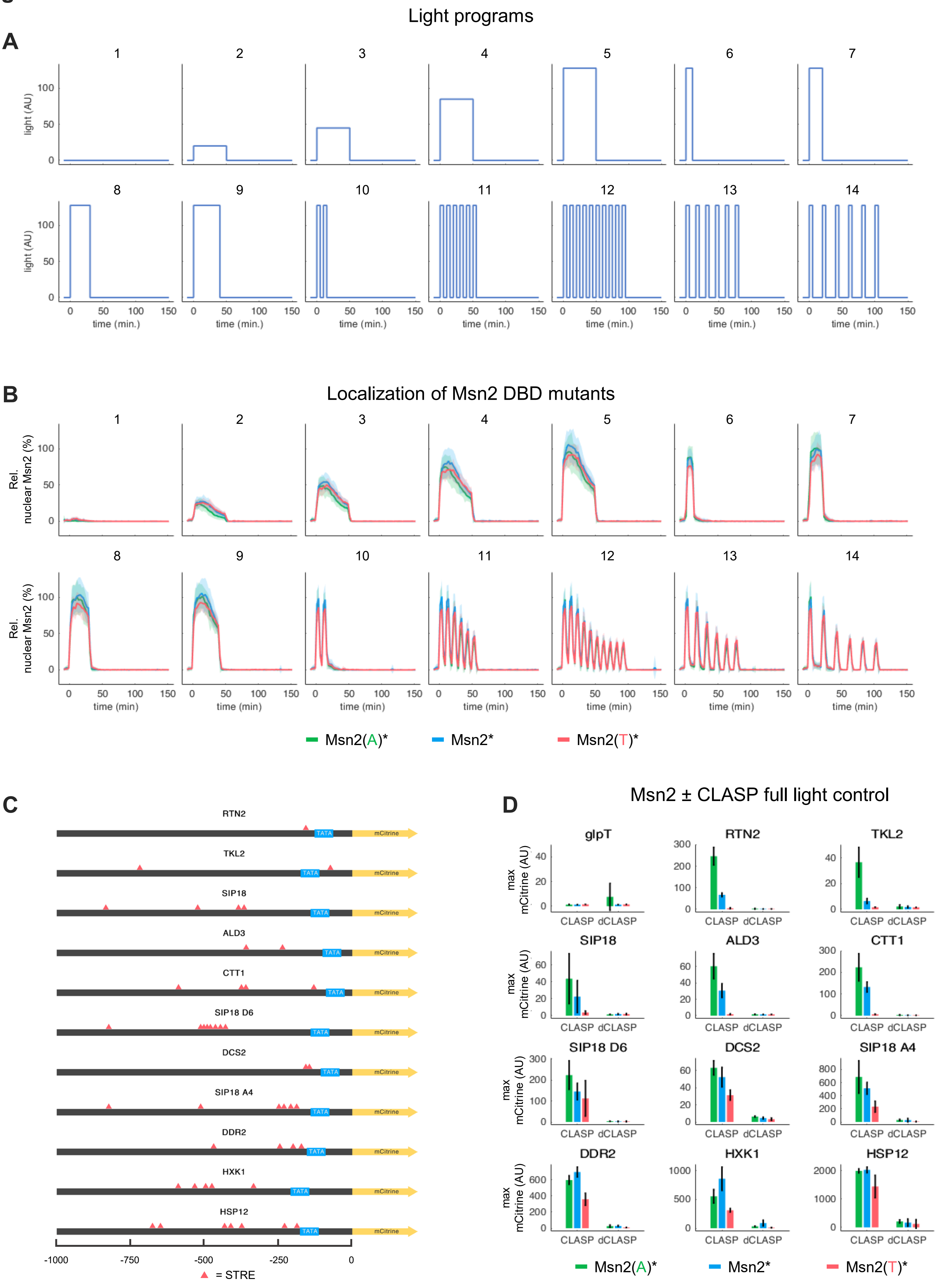

Light programs
A
1 2 3 4 5 6 7
8 9 10 11 12 13 14
light (AU)
time (min.)
B
Localization of Msn2 DBD mutants
Rel. nuclear Msn2 (%)
time (min)
Msn2(A)*
Msn2*
Msn2(T)*
C
RTN2
TKL2
SIP18
ALD3
CTT1
SIP18 D6
DCS2
SIP18 A4
DDR2
HXK1
HSP12
TATA
mCitrine
-1000
-750
-500
-250
0
= STRE
D
Msn2 ± CLASP full light control
glpT
RTN2
TKL2
SIP18
ALD3
CTT1
SIP18 D6
DCS2
SIP18 A4
DDR2
HXK1
HSP12
max mCitrine (AU)
CLASP
dCLASP
Msn2(A)*
Msn2*
Msn2(T)*

#### Figure S6

**A**

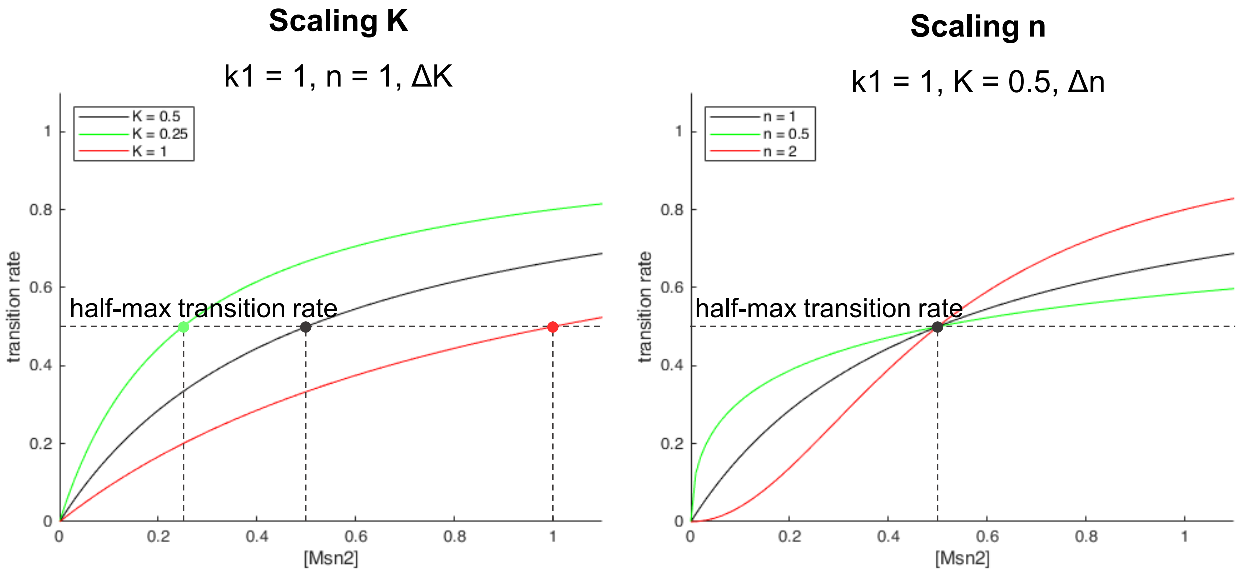


$$\text{transition rate} = \frac{k_1 Msn2(t)^n}{K^n + Msn2(t)^n}$$

**B**

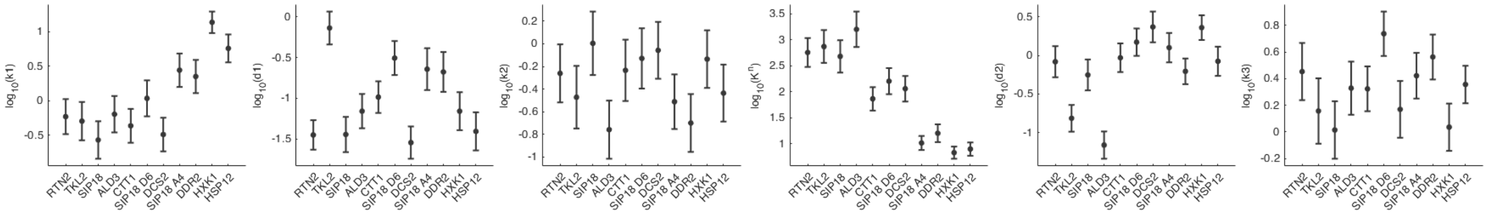


**C**

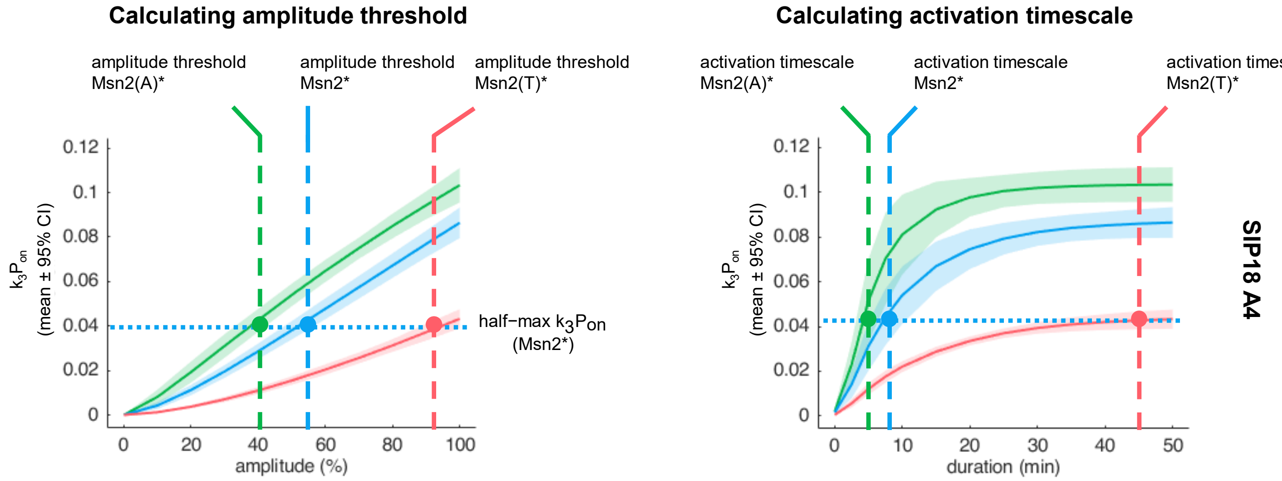

### Figure S7

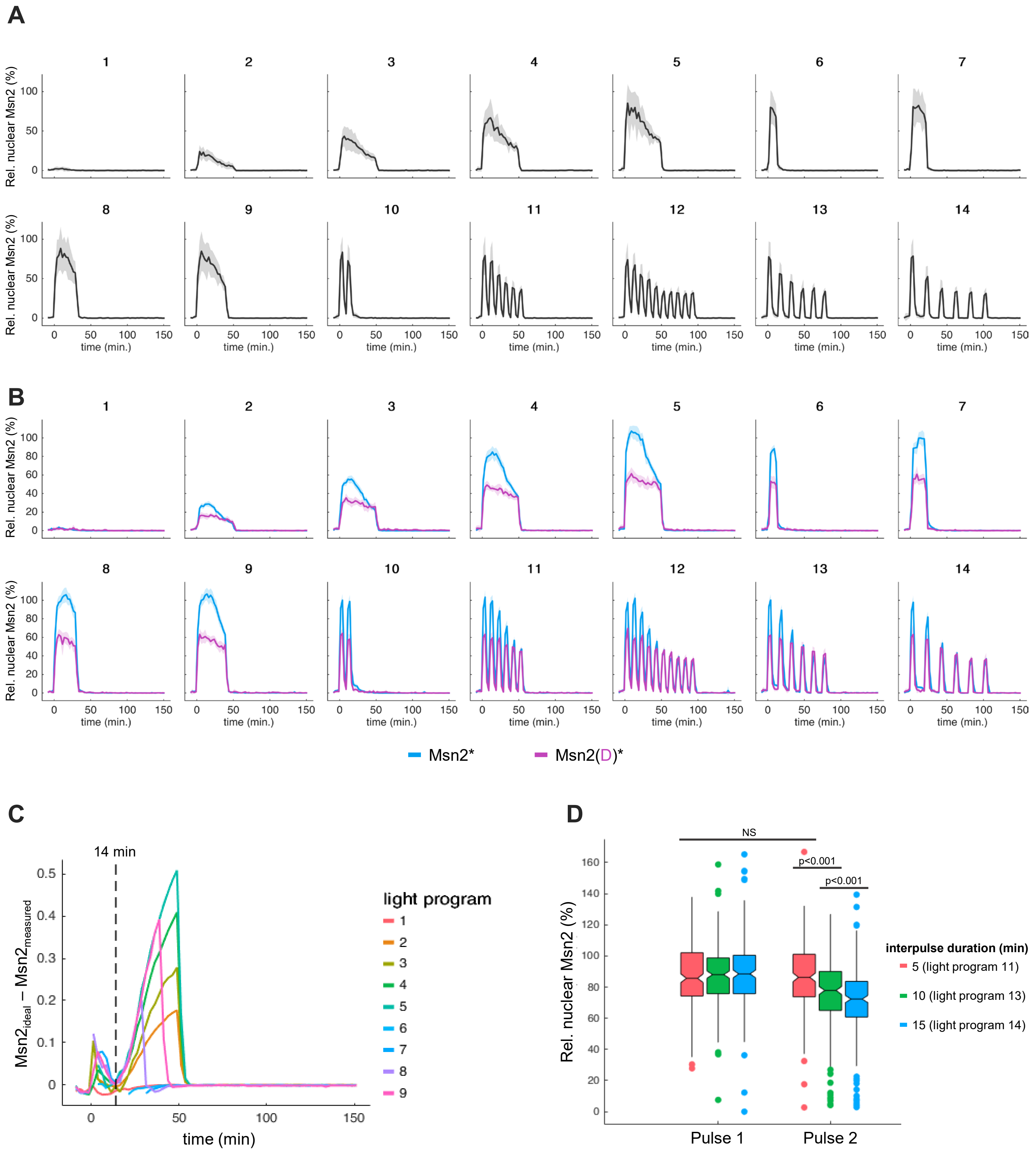

A
1
2
3
4
5
6
7
8
9
10
11
12
13
14
Rel. nuclear Msn2 (%)
time (min.)
B
Msn2*
Msn2(D)*
C
14 min
$Msn2_{ideal} - Msn2_{measured}$
time (min)
light program
D
NS
p<0.001
p<0.001
Rel. nuclear Msn2 (%)
Pulse 1
Pulse 2
interpulse duration (min)
5 (light program 11)
10 (light program 13)
15 (light program 14)

**Figure S8**

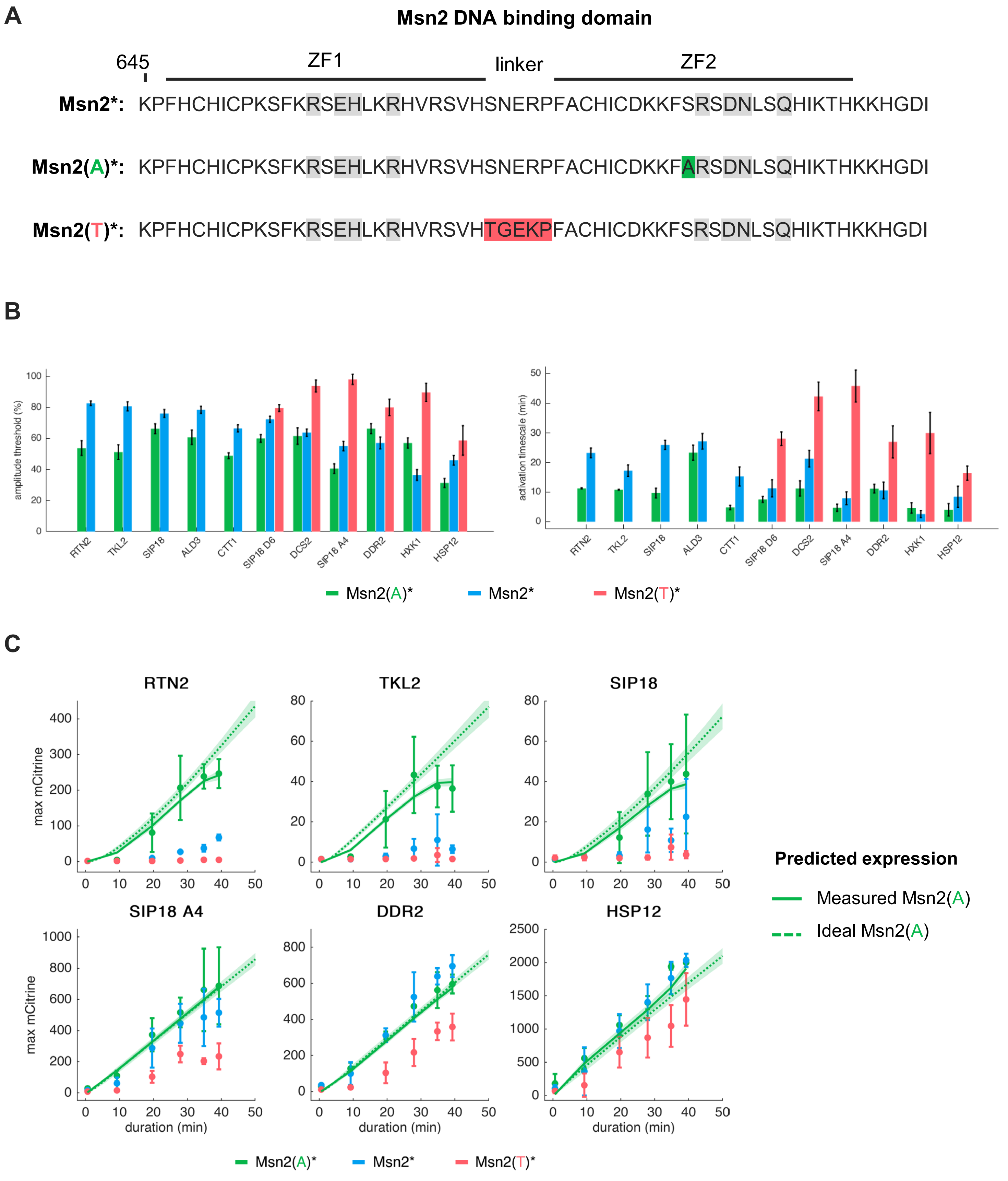
A
Msn2 DNA binding domain
645
ZF1
linker
ZF2
Msn2*: KPFHCHICPKSFKRSEHLKRHVRSVHSNERPFACHICDKKFSRSDNLSQHIKTHKKHGDI
Msn2(A)*: KPFHCHICPKSFKRSEHLKRHVRSVHSNERPFACHICDKKFARSDNLSQHIKTHKKHGDI
Msn2(T)*: KPFHCHICPKSFKRSEHLKRHVRSVHTGEKPFACHICDKKFSRSDNLSQHIKTHKKHGDI
B
amplitude threshold (%)
activation timescale (min)
RTN2
TKL2
SIP18
ALD3
CTT1
SIP18 D6
DCS2
SIP18 A4
DDR2
HXK1
HSP12
Msn2(A)*
Msn2*
Msn2(T)*
C
RTN2
TKL2
SIP18
SIP18 A4
DDR2
HSP12
max mCitrine
duration (min)
Predicted expression
Measured Msn2(A)
Ideal Msn2(A)
Msn2(A)*
Msn2*
Msn2(T)*

**Figure S9**

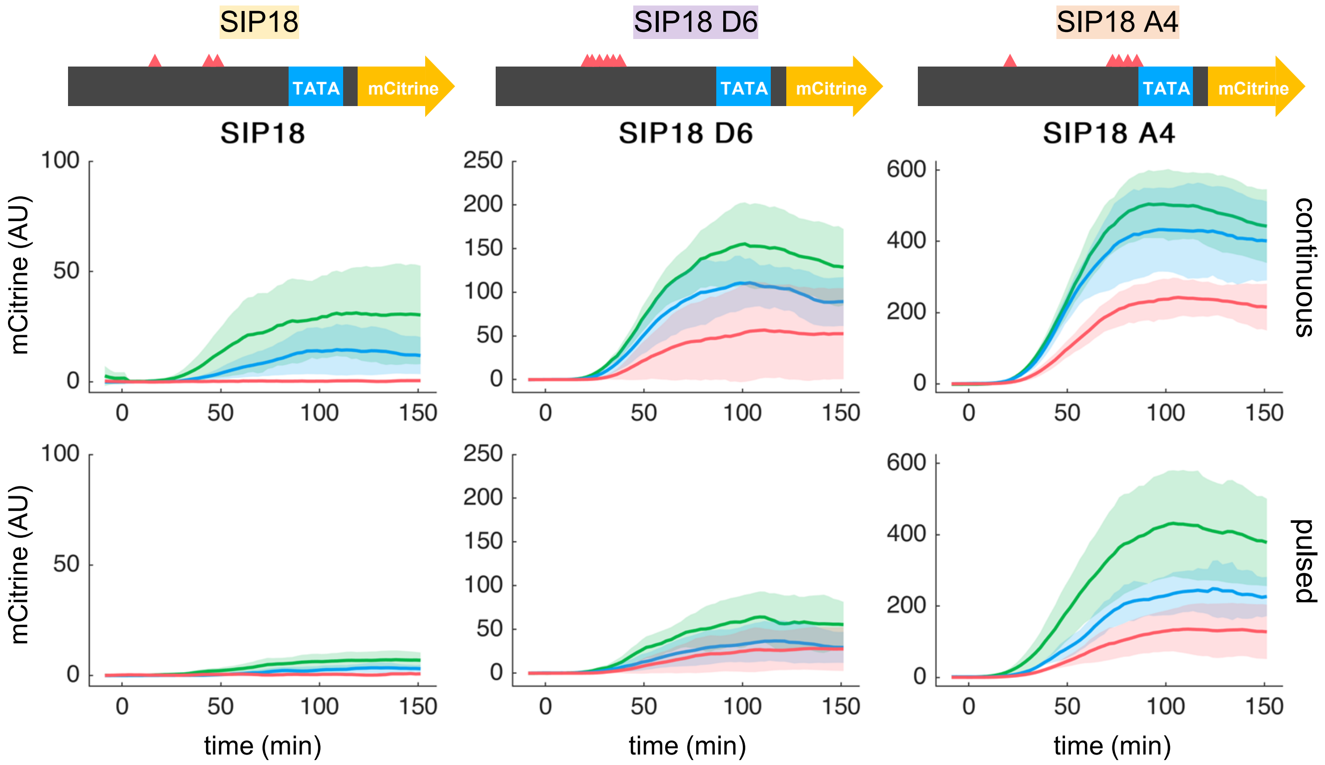
SIP18
SIP18 D6
SIP18 A4
TATA
mCitrine
mCitrine (AU)
time (min)
continuous
pulsed
0
50
100
150
200
250
400
600

### Figure S10

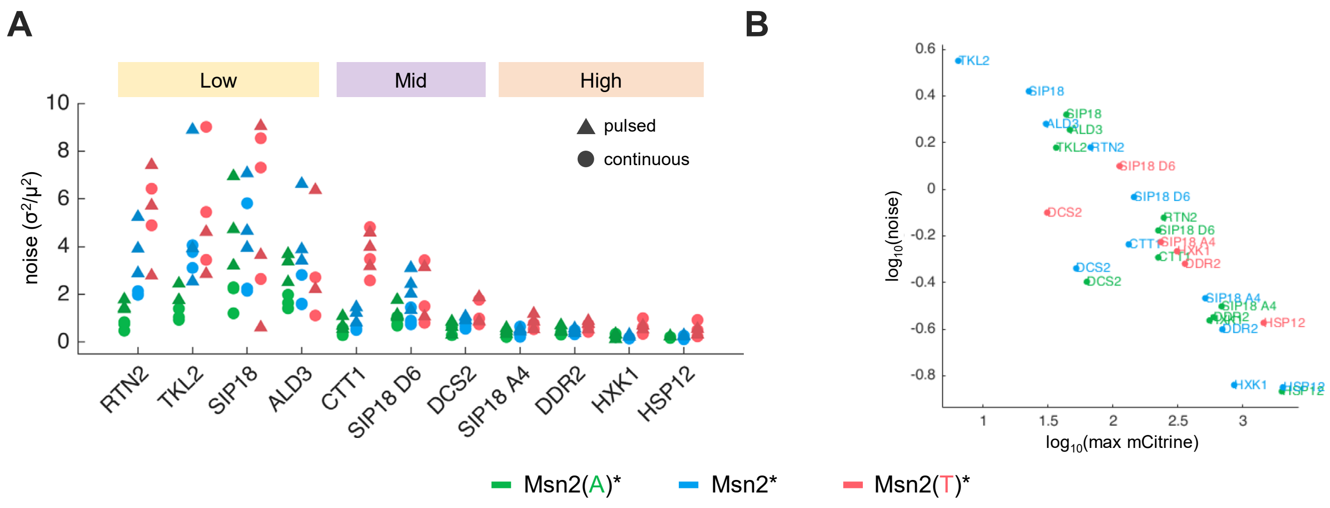
A
B
Low
Mid
High
pulsed
continuous
noise (σ²/μ²)
RTN2
TKL2
SIP18
ALD3
CTT1
SIP18 D6
DCS2
SIP18 A4
DDR2
HXK1
HSP12
log10(noise)
log10(max mCitrine)
Msn2(A)*
Msn2*
Msn2(T)*

### Figure S11

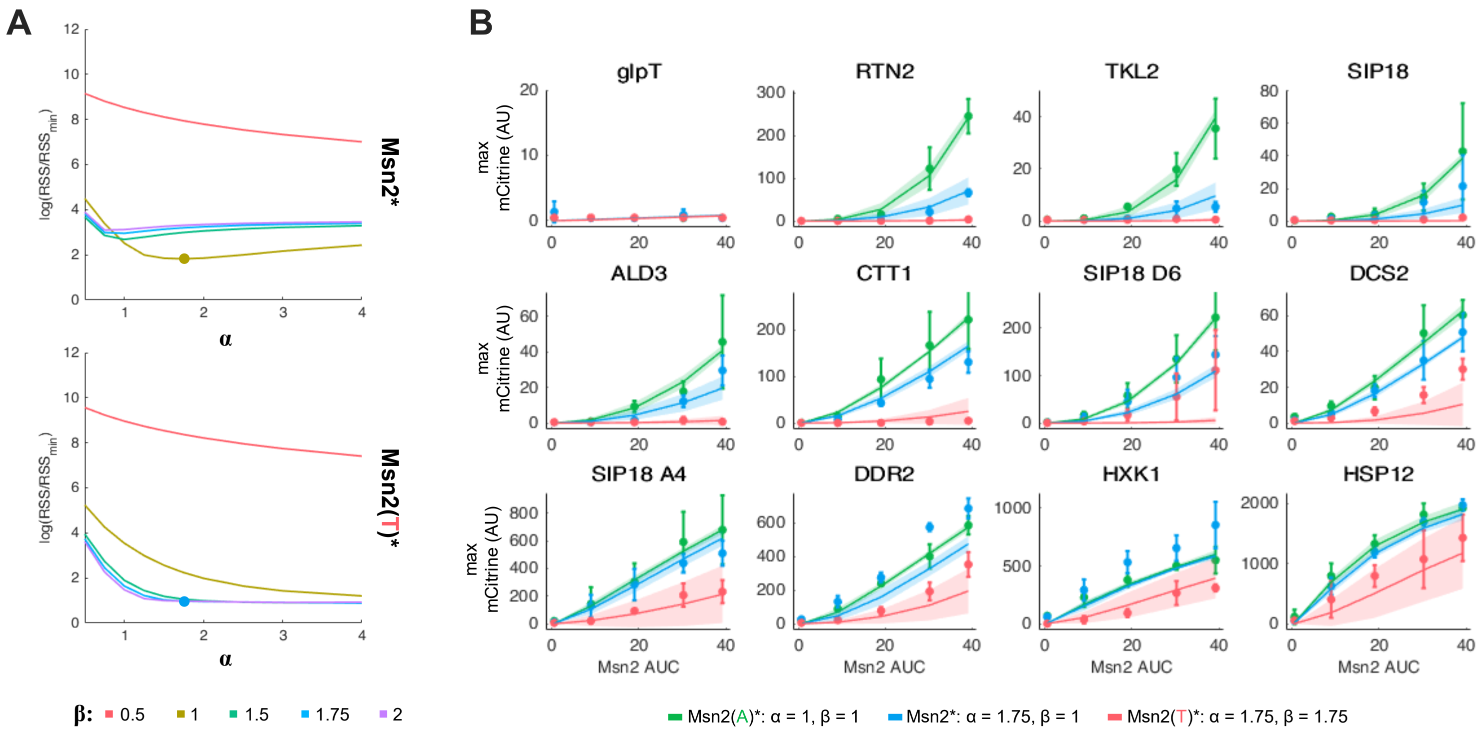
A
log(RSS/RSS$_{min}$)
α
Msn2*
Msn2(T)*
β: 0.5 1 1.5 1.75 2
B
glpT
RTN2
TKL2
SIP18
ALD3
CTT1
SIP18 D6
DCS2
SIP18 A4
DDR2
HXK1
HSP12
max mCitrine (AU)
Msn2 AUC
Msn2(A)*: α = 1, β = 1
Msn2*: α = 1.75, β = 1
Msn2(T)*: α = 1.75, β = 1.75

#### Figure S12

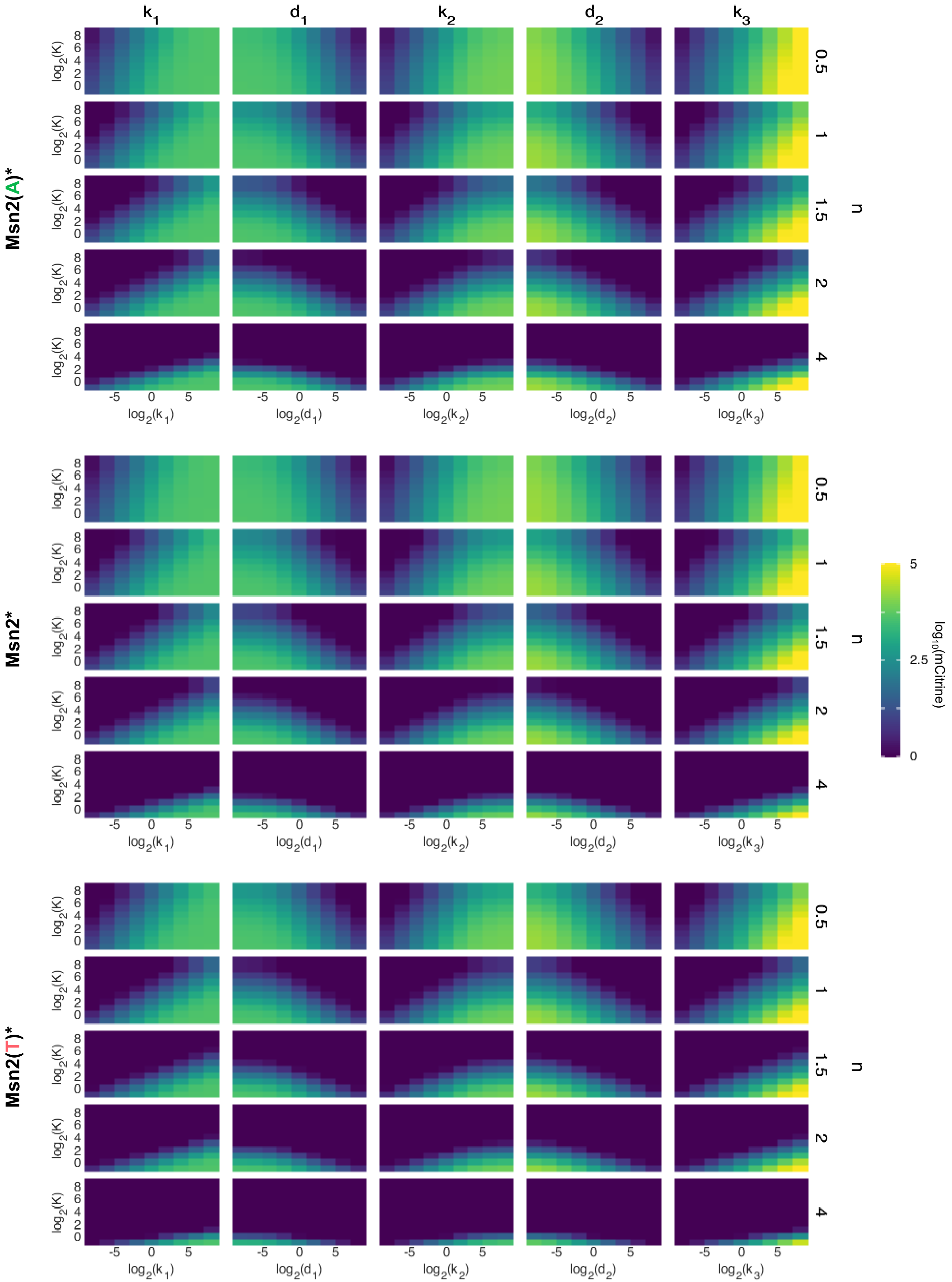

k1
d1
k2
d2
k3
Msn2(A)*
Msn2*
Msn2(T)*
log2(K)
log2(k1)
log2(d1)
log2(k2)
log2(d2)
log2(k3)
n
0.5
1
1.5
2
4
log10(mCitrine)
5
2.5
0

### Figure S13

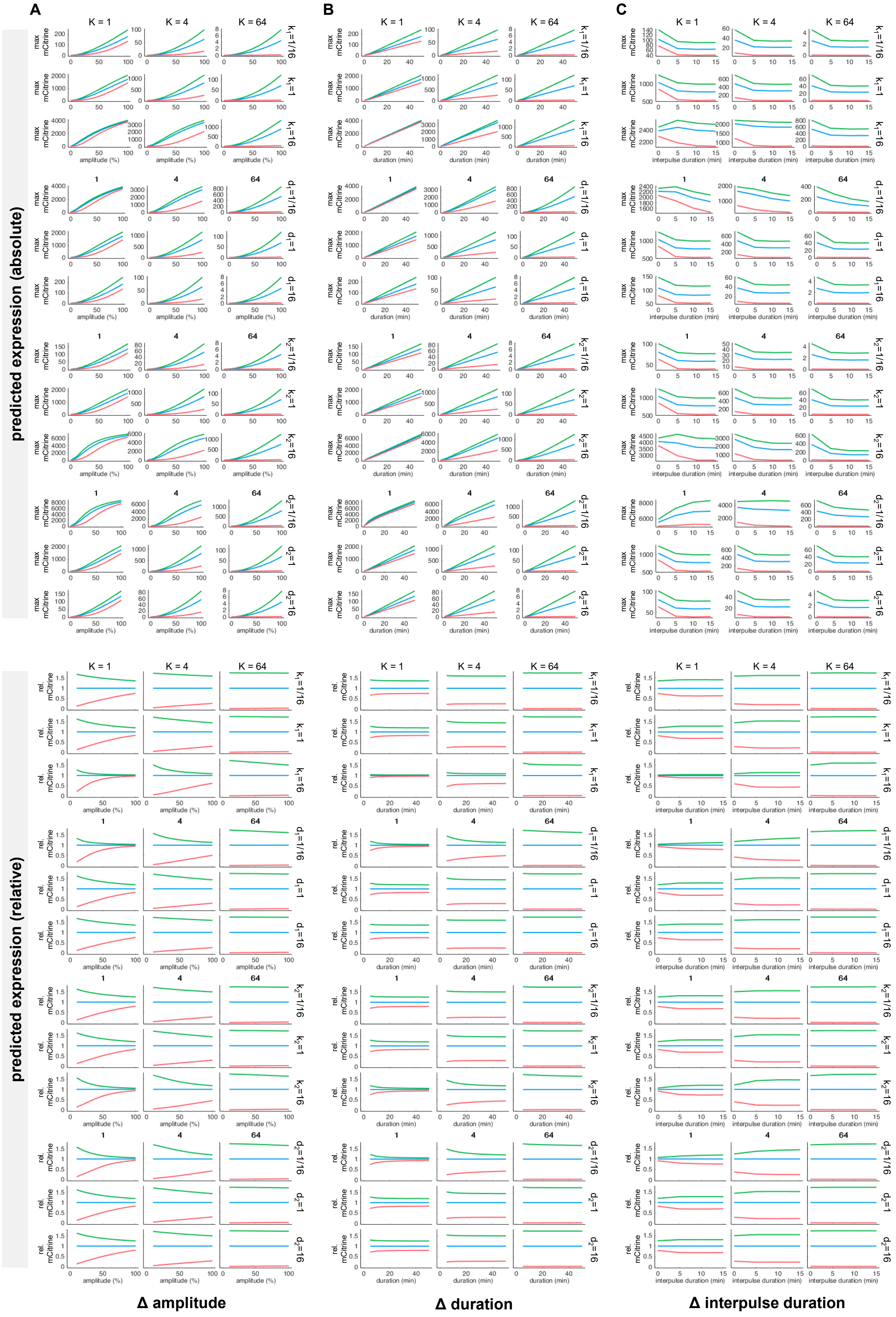
A
B
C
K = 1
K = 4
K = 64
predicted expression (absolute)
predicted expression (relative)
max mCitrine
rel. mCitrine
amplitude (%)
duration (min)
interpulse duration (min)
$k_1$=1/16
$k_1$=1
$k_1$=16
$d_1$=1/16
$d_1$=1
$d_1$=16
$k_2$=1/16
$k_2$=1
$k_2$=16
$d_2$=1/16
$d_2$=1
$d_2$=16
Δ amplitude
Δ duration
Δ interpulse duration

### Figure S14

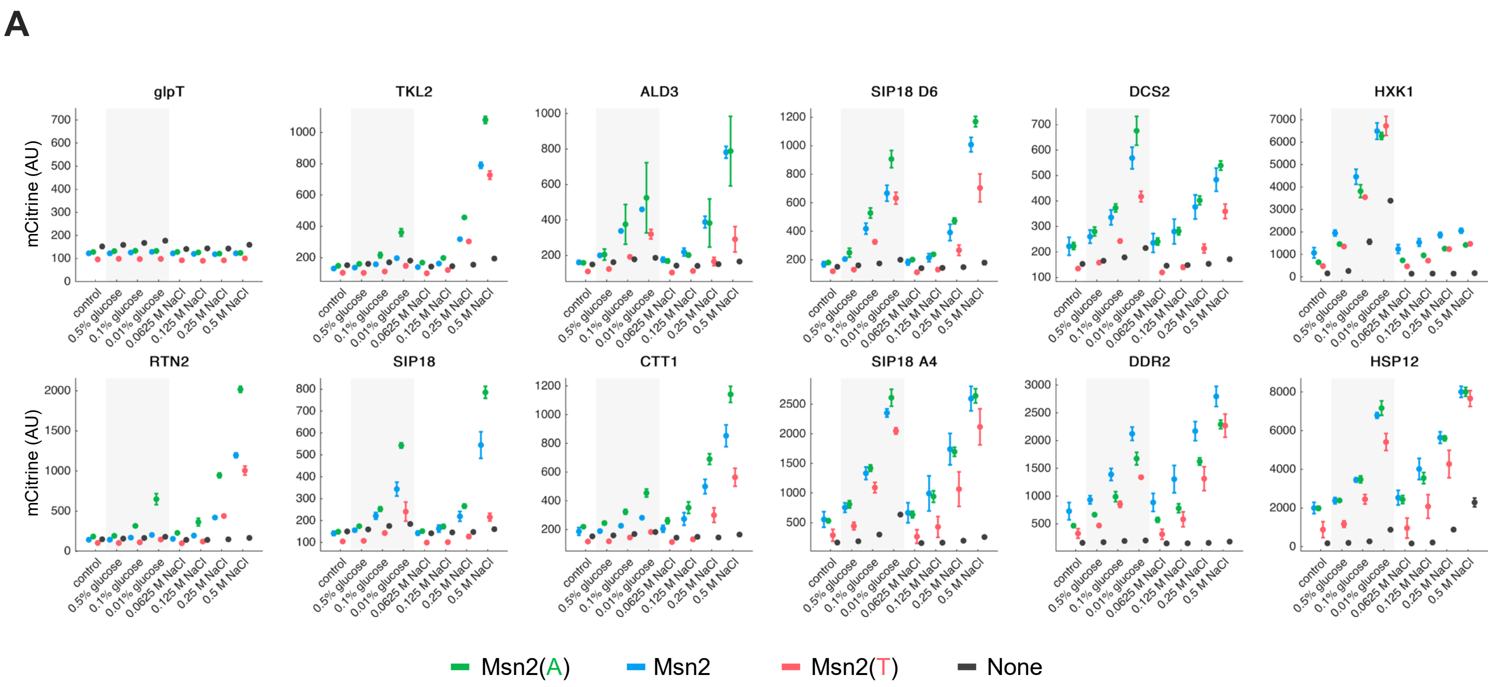

A
glpT
TKL2
ALD3
SIP18 D6
DCS2
HXK1
RTN2
SIP18
CTT1
SIP18 A4
DDR2
HSP12
mCitrine (AU)
control
0.5% glucose
0.1% glucose
0.01% glucose
0.0625 M NaCl
0.125 M NaCl
0.25 M NaCl
0.5 M NaCl
Msn2(A)
Msn2
Msn2(T)
None


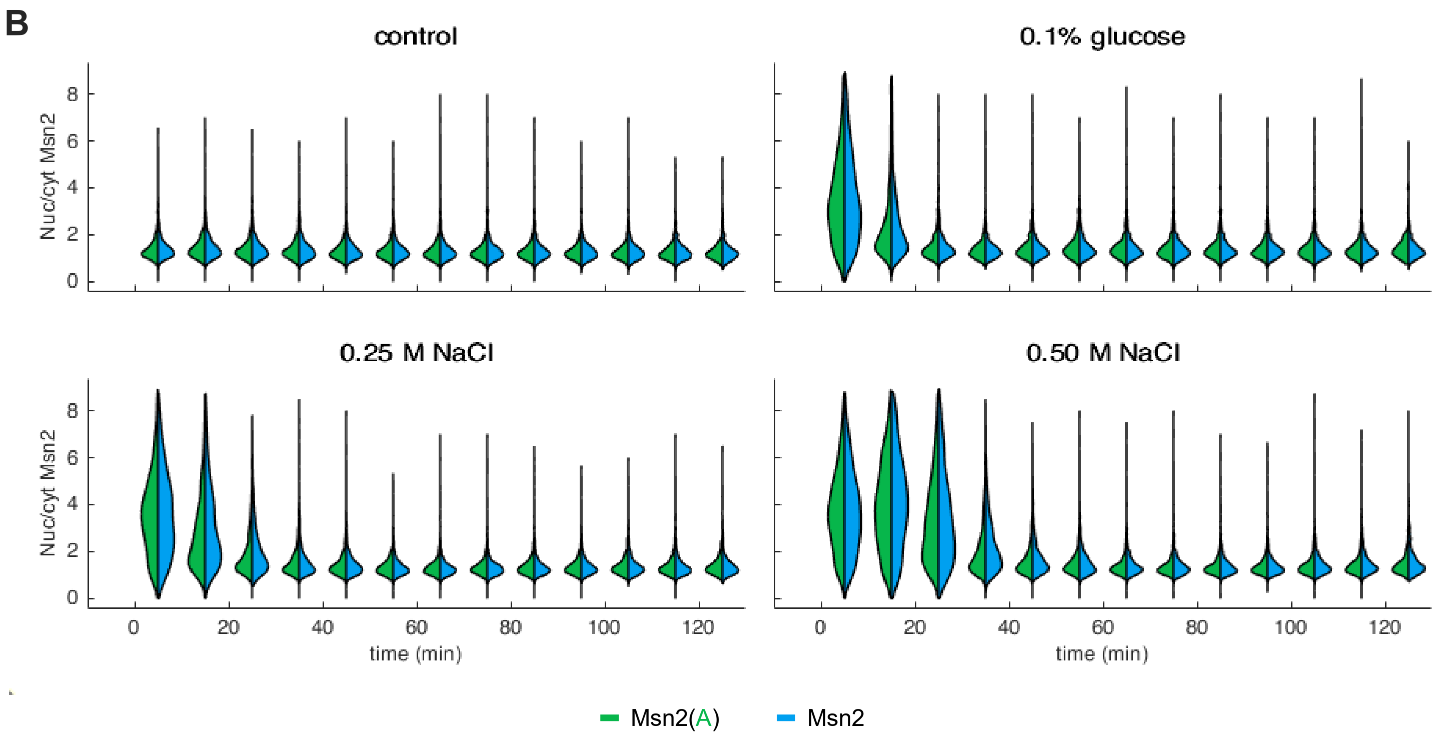

B
control
0.1% glucose
0.25 M NaCl
0.50 M NaCl
Nuc/cyt Msn2
time (min)
Msn2(A)
Msn2
